# PD-1 is induced on tumor-associated macrophages in obesity to directly restrain anti-tumor immunity

**DOI:** 10.1101/2022.11.12.515348

**Authors:** Jackie E. Bader, Melissa M. Wolf, Matthew Z. Madden, Bradley I. Reinfeld, Emily N. Arner, Emma S. Hathaway, KayLee K. Steiner, Gabriel A. Needle, Madelyn D. Landis, Matthew A. Cottam, Xiang Ye, Anthos Christofides, Vassiliki A. Boussiotis, Scott M. Haake, Kathryn E. Beckermann, W. Kimryn Rathmell, Alyssa H. Hasty, Jeffrey C. Rathmell

## Abstract

Obesity is a leading risk factor for progression and metastasis of many cancers^1,2^, yet can also promote improved survival for some cancers^3-5^ and enhance responses to some immune checkpoint blockade therapies^6-8^. The role of the immune system in the obesity-cancer connection and how obesity influences immunotherapy, however, remain unclear. While PD-1 expression by macrophages has been described^9-12^, we found that obesity selectively induced PD-1 on macrophages and that PD-1 directly impaired macrophage function. Single cell RNA sequencing of murine colorectal carcinoma tumors showed obesity remodeled myeloid and T cell populations, with fewer clonally expanded effector T cells and increased abundance of PD-1^+^ tumor-associated macrophages (TAM). Cytokines and molecules associated with obesity, including IL-6, leptin, and insulin, and the unsaturated fatty acid palmitate, induced PD-1 expression on macrophages in a glycolysis-dependent manner. PD-1^+^ TAMs had increased mitochondrial respiration and expression of genes regulating oxidative phosphorylation, lipid uptake and cell cycle while PD-1^-^ TAMs showed greater signatures of phagocytosis and antigen presentation to T cells. These patterns were directly regulated by PD-1, as recombinant PD-L1 reduced macrophage glycolysis and phagocytic capacity, and this was reversed with blocking PD-1 antibody. Conversely, PD-1-deficient *Pdcd1*^*-/-*^ TAMs had high rates of glycolysis, phagocytosis, and expression of MHC-II. Myeloid-specific PD-1 deficiency correlated with slower tumor growth, enhanced TAM antigen presentation capability, and increased CD8 T cell activation together with reduced markers of exhaustion. These findings show metabolic signaling in obesity induces PD-1-mediated suppression of TAM function and reveal a unique macrophage-specific mechanism to modulate immune tumor surveillance and checkpoint blockade. This may contribute to increased cancer risk yet improved response to PD-1 blockade in TAM-enriched tumors and obesity.

---

Over thirteen types of cancer have been associated with obesity, which is second only to smoking as a preventable cancer risk factor^13-15^ and becoming increasing prevalent worldwide. Conversely, obesity can prime for robust responses to immunotherapy through PD-1 blockade, in what has been termed the obesity paradox^16-20^. Though some obese patient cohorts have shown conflicting response to immune therapy ^21^, this is complicated by the use of body mass index (BMI) as a simplified and often unreliable indicator of health assessment and absence of considerations of other quantifiable features of metabolic health^22^. Studies investigating the effects of obesity on immunotherapy efficacy have largely focused on T cell phenotyping and reported that obesity can result in reduced tumor infiltrating T cells with a more naïve phenotype for those T cells that remain^23,24^. However, macrophages are well known to infiltrate and predominate tumor and adipose tissue microenvironments and act as key drivers of obesity-associated comorbidities including chronic inflammation, insulin resistance and lipid dysregulation^25^. Given the established link of macrophages in obesity and the abundance of macrophages in many tumor microenvironments, we sought to explore the role of TAM in obesity-associated cancer and response to immunotherapy.

## Obesity impairs tumor immune function

Diet induced obesity (DIO) can alter the tumor immune landscape^24^. A syngeneic subcutaneous tumor model using colorectal adenocarcinoma cell line, MC38, was employed here to test the effects of diet-induced obesity on tumor growth and the efficacy of anti-PD-1 immune checkpoint blockade (**Figure 1A**). Mice were given control low fat diet (LFD, 10% kcal fat) or high fat diet (HFD, 45% kcal fat), with both groups fed ad libitum for 20 weeks starting at 6 weeks of age. HFD mice gained significantly more body weight (**Extended Data Figure 1A**) and exhibited systemic obesity-associated metabolic changes including hyperglycemia and hyperinsulinemia, without significant changes in food intake (**Extended Data Figures 1 B-D**). After acclimation to diet feeding, mice were subcutaneously injected with syngeneic MC38 colorectal adenocarcinoma cells which were allowed to grow for 19 days. Consistent with obesity promoting cancer growth^24,26^, tumors grew more rapidly in mice fed HFD compared to LFD. After tumors were palpable, mice from both diets were treated with either anti-PD-1 or an IgG isotype control. Anti-PD-1 treatment appeared to attenuate tumor growth in both diet groups, however a significant decrease in tumor sizes only occurred in HFD-fed mice (**Figure 1B, Extended Data Figure 1E-J**). These data support the paradox that HFD can enhance both tumor growth and response to anti-PD-1.

**Figure 1.**
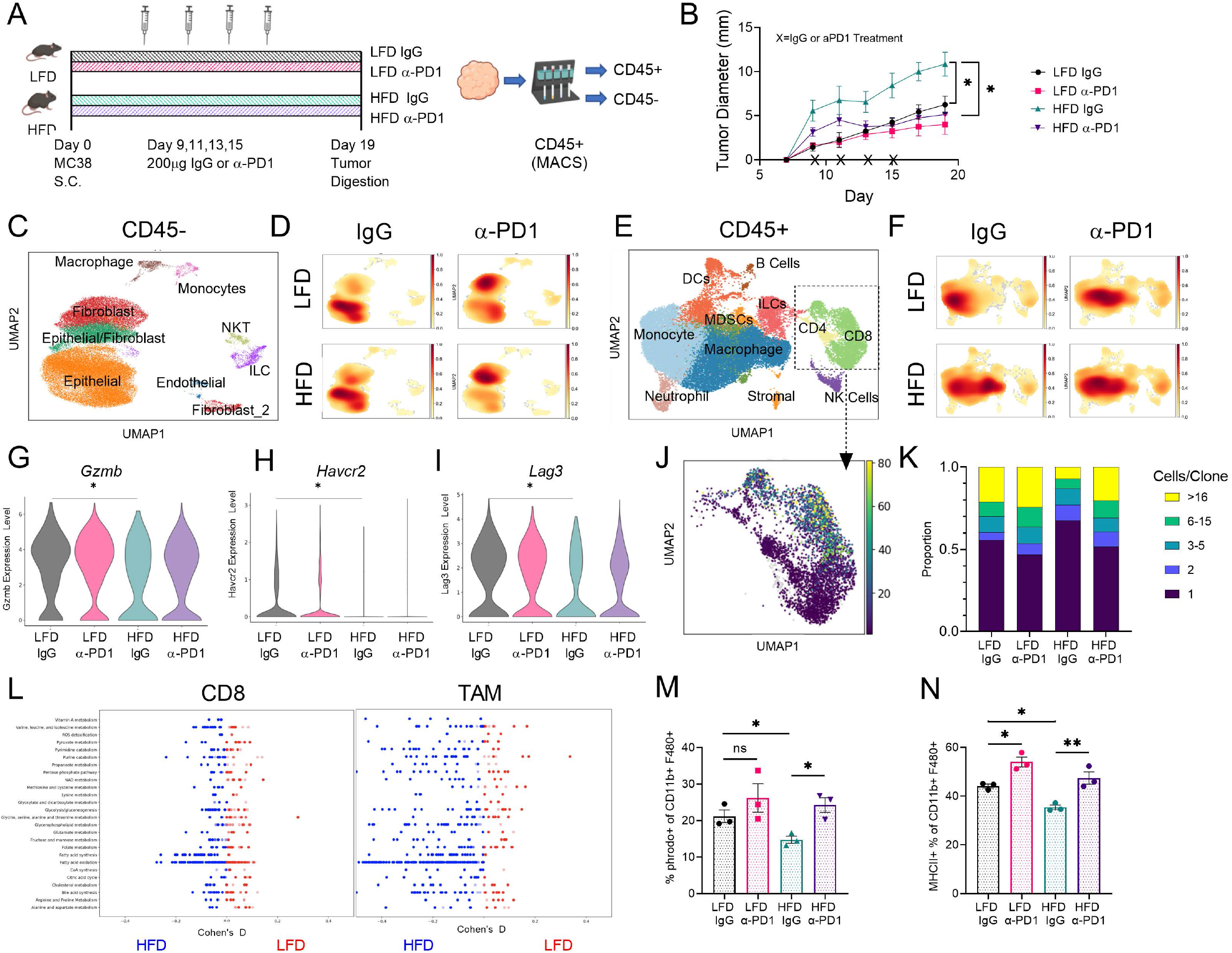
Obesity impairs anti-tumor immune cell function and metabolic signature. A) Schematic illustrating experimental approach. B) Growth curves of tumors in mice receiving IgG or anti-PD-1 treatment. C) Uniform Manifold Approximation and Projection (UMAP) embeddings of scRNAseq profiles from CD45^-^ cells from MC38 tumors annotated using SingleR. D) Density UMAPs of CD45^-^ cells from the 4 treatment groups. E) UMAP of CD45^+^ cells from MC38 tumors annotated using SingleR. F) Density UMAPs of CD45^+^ cells from the 4 treatment groups. G-I) Violin plot of gene expression within CD8 T cells. J) Combined UMAP of clone abundance from V(D)J sequencing of T cell clusters with productive TCRs. K) Proportion of total T cells with paired TCR-a and TCR-b sequences. L) Compass analysis of scRNAseq data represented as differential activity of metabolic reactions from CD8 and Macrophage identified clusters comparing LFD IgG to HFD IgG groups partitioned by Recon2 pathways. M) Percent of phrodo^+^ cells gated on live CD11b^+^ F480^+^ TAMs from MC38 tumors. N) Percent of MHCII^+^ TAMs from MC38 tumors. For scRNA sequencing in C-L, data are representative of one experiment, n = 3 pooled tumors from LFD mice and n = 3 pooled tumors from HFD mice. For B, M, and N each data point represents a biological replicate; data are mean ± s.e.m. Data from B, M, and N represent ≥ three independent experiments with ≥ 3 mice per group. P values were calculated using a One-way ANOVA (B) or Wilcoxon rank-sum test (G-I) or an unpaired two-tailed t-test (m-n). ns=p>0.05; * p≤0.05; ** p≤0.01.

To assess the effects of diet and immunotherapy at cellular detail, fresh CD45^+^ and CD45^-^ cells were isolated from tumors and subjected to single cell RNA sequencing (scRNAseq). Despite significant differences in tumor size, diet alone did not lead to a striking change in the relative abundance of CD45^-^ epithelial cells and fibroblasts (**Figure 1C-D, Extended Data Figure 2A**). Anti-PD-1 treatment, however, resulted in a shift from epithelial to fibroblast-identified cells that may reflect the elimination of tumor cells. The CD45^+^ fraction, in contrast, revealed significant alterations to the tumor immune cell landscape in the HFD group **(Figure 1E-F, Extended Data Figure 2B-C)**. HFD was sufficient to reduce the frequency of tumor infiltrating effector CD8 T cells (TIL), which was rescued with anti-PD-1 treatment. CD8 TIL from HFD treated mice exhibited decreased effector molecule expression with reduced *Gzmb*, and less exhausted phenotypes with reduced *Havcr2* (TIM3) and *Lag3* coinhibitory receptor expression that were reversed upon anti-PD-1 treatment (**Figure 1G-I, Extended Data Figure 2D, 3A-E**). TCR sequencing revealed that CD8 T cell clonality, a measure of the number of expanded clones and an indicator of anti-tumor activity, was decreased in the HFD group (**Figure 1J-K, Extended Data Figure 2E-F**). Interestingly, anti-PD-1 treatment increased CD8 cell clonality to a similar proportion between LFD and HFD groups despite HFD starting at a reduced clonal state.

More evident than shifts in T cell populations, however, were changes in the frequencies and subsets of TAM in HFD or anti-PD-1 treatment (**Figure 1E-F, Extended Data Figure 2B**). In contrast to a decrease in CD8 T cells in HFD, tumor infiltrating macrophages were sharply increased with obesity. We applied Compass to the scRNAseq gene expression to better assess metabolic states of CD8 T cells and TAMs with LFD vs HFD ^27^. CD8 T cells had a moderately altered metabolic state in response to HFD treatment, with increased fatty acid synthesis and FAO pathways most notable (**Figure 1L, Extended Data Figure 2G**). This increase was also observed in TAMs in response to HFD but to a greater magnitude than T cells and TAM had broadly altered metabolism in HFD treated mice. Importantly, TAM anti-tumor function including phagocytosis and antigen presentation via MHCII expression were significantly decreased in HFD and appeared to be rescued upon anti-PD-1 treatment (**Figure 1 M-N**). HFD TAMs also preferentially utilized glycolysis compared to LFD TAMs and this was increased with anti-PD-1 treatment (**Extended Data Figure 3F-H**). HFD thus results in fewer and less activated and clonally expanded CD8 T cells while reprogramming metabolism and impairing phagocytosis and antigen presentation of TAM and that these phenotypes are reversed upon anti-PD-1 treatment.

## Obesity selectively induces TAM to express PD-1

Obesity-associated immunotherapy responses may occur through changes to T cell activation or TAM sensitivity to anti-PD-1. Consistent with literature, TAMs expressed detectable, yet low levels of PD-1 compared to CD8 T cells in humans (**Figure 2A, Extended Data Figure 4A-B**). Analysis of a human scRNA-seq database showed PD-1 was expressed in myeloid cells of many cancer types (**Extended Date Figure 4C-E**). Similarly, TAMs were more frequently PD-1^+^ than macrophages in normal adjacent tissue in two independent scRNAseq datasets of colon cancer and kidney cancer patient samples ^28,29^ (**Figure 2B, Extended Data Figure 4G**). However, PD-1^+^ TAM frequencies did not correlate with tumor grade (**Extended Data Figure 4F-H**). Immunohistochemistry staining on a tumor microarray of over 100 resected tumors from patients with kidney cancer, revealed a clear co-localization of PD-1 on CD163^+^ myeloid cells in addition to PD-1 colocalized with CD8 T cells (**Figure 2C**). PD-1^+^ CD163^+^ myeloid and CD8 T cells were positively correlated to suggest that increased exhausted T cells are associated with increased PD-1 expressing macrophages (**Figure 2D**). The frequency of CD68^+^ TAMs showed a trend for increased abundance in overweight and obese patients compared to lean patients (**Figure 2E**). Analysis of differentially expressed genes in these human PD-1^+^ TAMs compared to PD-1^-^ TAMs, revealed increased genes associated with the ribosome and corona virus KEGG pathways in PD-1^-^ TAMs and increased insulin secretion KEGG pathway genes for PD-1^+^ TAMS suggesting a greater anabolic and inflammatory state of PD-1^-^ TAM relative to an insulin response in PD-1^+^ TAM (**Extended Data Figure 4I**).

**Figure 2.**
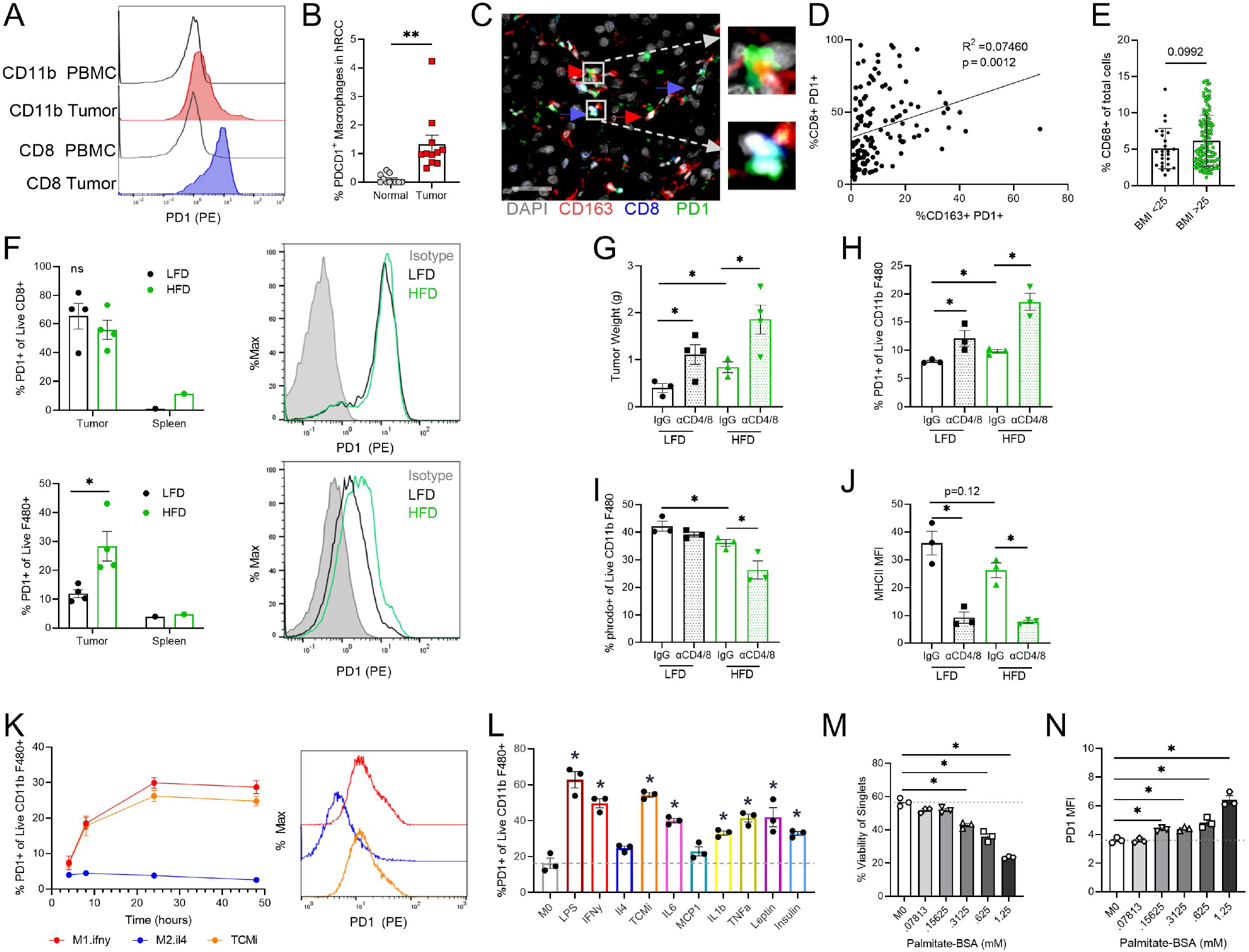
Obesity and obesity associated signaling increased macrophage specific PD-1. A) Representative histogram of PD-1 expression on human peripheral blood mononuclear cells (PBMCs) and matched ccRCC tumor in CD11b^+^ TAMs and CD8^+^ TILs. B) Percent *PDCD1*^+^ TAMs compared to adjacent normal tissue from human ccRCC samples measured using gene expression from scRNAseq (n=11patients). C) Representative image of Tumor Microarray (TMA) staining identifying PD-1 colocalizing with CD163^+^ and CD8^+^ cells. Scale bar 50mm. D) Correlation of PD-1^+^ CD8 cells vs PD-1^+^ CD163 cells from TMA staining (n=163). E) Percentage of CD68^+^ macrophages in TMA staining with patient clinical data categorized as lean or overweight (lean BMI < 25; overweight BMI ≥ 25) (n=25 leanpatients; n=135 overwight/obese). F) Percent PD-1^+^ within CD8^+^ and F480^+^ gated live cells from LFD and HFD treated mice with representative histogram. G-J) Tumor weights, PD-1+ expression, % phrodo^+^ phagocytic cells, and MHCII expression on CD11b+/F480+ cells on day 14 of MC38 tumors following aCD4 + aCD8 T cell depletion in mice fed LFD or HFD for 20 weeks (n=3-4 mice). K) PD-1 expression of BMDMs following 4-, 8-, 24- and 48-hour exposure to IFNg, IL-4 or tumor-conditioned media (TCM). L) PD-1 expression of BMDMs following 24-hour exposure to LPS (1ng/mL), IFNg (50ng/mL), IL-4 (50ng/mL), TCM (25% conditioned media), IL-6 (50ng/mL), MCP-1 (50ng/mL), IL-1β (50ng/mL), TNF-α (50ng/mL), leptin (5μg/mL), or insulin (100mg/mL) (n=3 mice). M-N) Viability and PD-1 expression of BMDM following 24-hour treatment with increasing doses of BSA-conjugated palmitic acid (n=3 mice). Each data point represents a biological replicate presented as mean ± s.e.m.; Data in F-J represent ≥ three independent experiments with ≥ 3 mice per group. P values were calculated using an unpaired two-tailed t-test (F) or a one-way ANOVA compared to IgG or M0 (unstimulated BMDM) (G-M). ns p>0.05, *p≤0.05).

Like human, macrophages and myeloid cells in the murine DIO tumor model expressed PD-1, albeit at lower levels than CD8 and CD4 T cells (**Extended Data Figure 5A**). As expected, myeloid cells had the highest *Cd279* (PD-L1) expression **(Extended Data Figure 5B)**. Importantly, HFD significantly increased PD-1 expressing TAMs but did not increase PD-1 on splenic macrophages or in exhausted T cells (**Figure 2F, Extended Data Figure 5C**). Macrophage-specific PD-1 did not change in response to an acute HFD treatment of 2 weeks (**Extended Data Figure 5D-E**), but instead developed at later time points concordant with onset of metabolic dysfunction. To determine the extent of T cell influence on tumor growth and TAM PD-1 expression, T cells were depleted by treatment with anti-CD4 and anti-CD8 antibodies. As expected, this enhanced tumor growth, with an additive effect in the HFD group (**Figure 2G, Extended Data Figure 5F-H**). PD-1 expressing TAMs were also further increased (**Figure 2H**) to demonstrate macrophage PD-1 did not rely on but instead was suppressed by T cells. TAM specific PD-1 expression was also associated with macrophage dysfunction, demonstrated by decreased phagocytosis and MHCII expression (**Figure 2 I-J**). Although MHCII expression does not directly associate to CD8 cells, it aids indirectly in the presentation to other immune cells. These data support a model in which obesity and metabolic dysfunction directly drive macrophage PD-1 expression independent of T cell signaling.

The regulation of macrophage PD-1 expression was next assessed. Bone marrow derived macrophages (BMDMs) were stimulated to M1 and M2 polarization conditions with IFNγ or IL4 treatments, respectively. To model TAM, BMDMs were conditioned with MC38 tumor conditioned media (TCM). Both M1-like and TCM groups showed increased PD-1 expression as early as 8 hours following stimulation, while PD-1 expression did not change on M2-like macrophages (**Figure 2K**). Expression of PD-L1 increased similarly to PD-1 in M1 and TCM groups (**Extended Data Figure 5I**). To further explore cytokine influence on macrophage PD-1 response we treated BMDM with cytokines and adipokines which are often increased in obesity^30,31^. The pro-inflammatory and obesity-associated cytokines and adipokines including IL6, IL1β, TNFα, leptin, and insulin all increased PD-1^+^ on BMDMs (**Figure 2L, Extended Data Figure 5J**). PD-L1, the ligand for PD-1 most notably expressed on cancer cells and TAMs^32^, also increased upon treatment with LPS, IFNγ, TCM, TNFα and Leptin (**Extended Data Figure 5K**). Obesity also increases circulating free fatty acids^33^ and treatment with unsaturated fatty acid, palmitic acid C16:0, resulted in a dose response toxicity which inversely correlated with an increase in PD-1 expression (**Figure 2M-N**). While previously established that TAMs can express PD-1, which negatively correlates with phagocytic function^9,34,35^, our data now show HFD and obesity induces TAM PD-1 expression through increased inflammatory cytokines, leptin, insulin, and free fatty acids.

## PD-1 directly impairs macrophage function and anabolic metabolism

To determine if PD-1 signaling affected macrophages, we examined tumors, purified macrophage cultures, and genetically targeted PD-1 *in vivo*. Differential gene expression patterns from PD-1^+^ to PD-1^-^ TAMs were identified from scRNA analysis (**Figure 1, 3A, Extended Data Figure 6, Extended Data Table 1**). PD-1^+^ expressing TAMs exhibited a differential gene expression in biological pathways associated with IL-4 response, oxidative phosphorylation, and cell cycle, while PD-1^-^ TAMs were enriched for biological pathways associated with phagocytosis, T cell activation and TNFα and IL6 production (**Figure 3A, Extended Data Figure 6**). Of note, among the differentially expressed genes in T cell activation *B2m* was enriched in PD1^-^ TAMs, supporting greater antigen presentation capacity of these macrophages to CD8 T cells. Consistent with these differentially expressed gene sets, sorted PD-1 high and PD-1 low tumor conditioned macrophages confirmed an increase in oxidative phosphorylation among the PD-1 high groups (**Figure 3B-C, Extended Data Figure 7A**). Additionally, gating on PD-1 hi and low tumor associated macrophages from MC38 tumors by flow cytometry affirmed that PD-1^+^ TAMs have reduced phagocytosis along with increase in lipid uptake (**Figure 3D-E, Extended Data Figure 7B**). These data are consistent with PD-1 as a regulator of macrophage metabolism and function.

**Figure 3.**
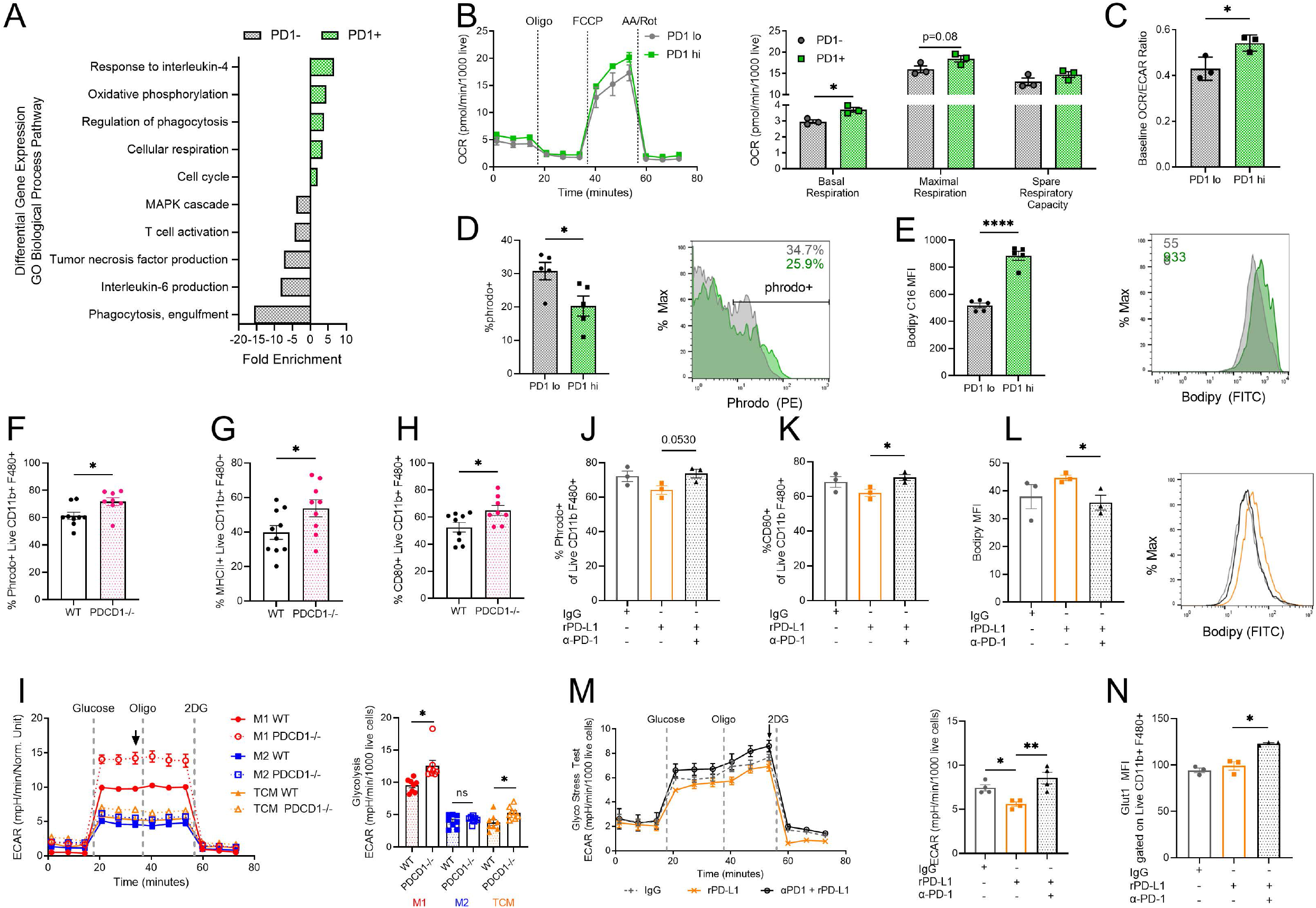
Anti-PD-1 directly increases macrophage anti-tumor function and metabolism. A) Enrichment of gene ontology biological processes scores of *Pdcd1*^-^ versus *Pdcd1*^+^ TAMs from MC38 tumor scRNAseq. B) Representative extracellular flux analysis oxygen consumption rate (OCR) of PD-1 hi and PD-1 low FACS-sorted TCM-treated BMDMs with quantified basal respiration, maximal respiration, and spare respiratory capacity (each dot represents technical replicates; data representative of two independent experiments). C) Basal OCR/ECAR ratio of PD-1 hi and PD-1 low FACS-sorted TCM-treated BMDMs (n=3 technical replicates). D) Percent phrodo^+^ cells gated on live CD11b^+^ F480^+^ PD-1 hi or PD-1 low macrophages from MC38 tumors (n=5 mice). E) Palmitate analog Bodipy C:16 F-H median fluorescence intensity and representative histograms in PD-1 hi and PD-1 low TAMs from MC38 tumors (n=5 mice) F-I) Percent of phrodo+, MHCII+, CD80+ peritoneal macrophages isolated from WT or *Pdcd1*^*-/-*^ male mice (n=8-9 mice). I) Representative extracellular flux analysis of BMDM from WT or *Pdcd1*^*-/-*^ mice stimulated for 24hours with IFN-g (M1), IL-4 (M2), or TCM and quantification of glycolysis (n=8-10 mice). J-L) Quantification by flow cytometry of % phrodo^+^, CD80^+^ and MFI of bodipy of BMDMs stimulated with TCM for 24hours followed by treatment with rmPD-L1 and/or anti-PD-1 for an additional 24hours. M) Representative extracellular flux analysis of BMDM with indicated injections of compounds following rmPD-L1 and anti-PD-1 treatment with quantified maximal glycolysis (n=4 mice). N) Flow cytometry quantification of GLUT1 MFI from BMDMs following respective treatments (n=3 mice). Data represent ≥ two independent experiments with ≥ 3 mice per group data are mean ± s.e.m. (ns p>0.05, *p≤0.05).

To directly test if PD-1 deficient macrophages exhibit enhanced function, MC38 tumors from *Pdcd1*^*-/-*^ mice were analyzed. Whole body PD-1 deficiency strongly suppressed MC38 syngeneic tumor growth. Isolating peritoneal macrophages from tumor-exposed mice, however, revealed that *Pdcd1*^*-/-*^ macrophages had a more pro-inflammatory state as shown by increased phagocytosis, MHCII and CD80 expression and increased glucose uptake (**Figure 3F-H, Extended Data Figure 8A**). Further, *in vitro* stimulated M1-like, M2-like and TCM BMDMs from *Pdcd1*^*-/-*^ mice preferentially increased glycolysis as indicated by extracellular acidification (**Figure 3I, Extended Data Figure 8B-C**). Glycolysis was most increased in M1 and TCM groups which exhibit the highest PD-1 expression. Conversely, PD-1 expression decreased *in vitro* following treatment with glucose antagonist, 2DG, in highly glycolytic macrophages but not following treatment with pan-glutamine antagonist, DON, or fatty acid oxidation inhibitor, etomoxir (**Extended Data Figure 8D-F**). Increased glycolysis correlates with and is thus essential for macrophage to induce expression of PD-1.

To test if PD-1^+^ macrophages directly respond to anti-PD-1 treatment, BMDMs were treated with recombinant PD-L1 or anti-PD-1 blocking antibody. Saturating PD-1 on macrophages with recombinant mouse PD-L1 (rmPD-L1) resulted in decreased phagocytosis, CD80 expression, and lipid uptake while co-treatment with a PD-1 blocking antibody restored and enhanced pro-inflammatory metabolism and function. PD-1 blockade increased phagocytosis and expression of co-stimulatory ligands (**Figure 3J-K**). In addition, blockade reduced lipid uptake while increasing glycolysis and expression of GLUT1 protein (**Figure 3L-N, Extended Data Figure 8G-J**). Taken together, these data indicate that PD-1 can directly alter macrophage metabolism and reduce macrophage ability to phagocytose targets and stimulate T cells.

## Macrophage specific PD-1 restricts anti-tumor immunity

Our data show obesity-induced PD-1 can directly impair macrophages. We next tested if PD-1 directly regulated macrophages in the tumor microenvironment or if PD-1 effects depended solely on T cells. T cell-depleted tumors had increased growth but still responded to anti-PD-1 treatment (**Figure 4A**). Interestingly, TAMs in T cell depleted tumor microenvironments responded to anti-PD-1 treatment through increased phagocytosis and glucose uptake and reduced lipid uptake (**Figure 4B-D, Extended Data Figure 9A-C**). Consistent with literature^36^, whole body PD-1 knockout mice had significantly diminished tumor growth (**Figure 4E**). Within these tumors, TAMs had increased GLUT1 expression and increased MHCII and CD86, suggesting increased glycolysis and antigen presentation (**Figure 4F-H**). Given the established impact of PD-1 depletion in T cells, it was unclear whether this enhanced antitumor phenotype in the TAMs was a direct result of PD-1 depletion or an indirect effect from sustained T cell activation in the PD-1 deficient mice. To test this, *LysMCre Pdcd1*^*fl/fl*^ animals were subjected to the same MC38 tumor model. PD-1-deficient myeloid cells effectively reduced tumor growth, consistent with the full body knockout, and previous results from Strauss, et. al.^11^ (**Figure 4I**). Importantly, PD-1 deficient TAMs exhibited enhanced phagocytosis, reduced lipid uptake, and broadly increased metabolic activity compared to WT TAMs suggesting increased anabolic metabolism and an improved ability to present antigen and activate T cells (**Figure 4J-L, Extended Data Figure 10A-B, E-F**). Consistent with increased T cell interactions, PD-1 deficient TAMs improved T-cell activation and proliferation when co-cultured with naïve non-tumor bearing splenic OVA specific CD8 cells and its antigen OVA peptide, SIINFEKL. (**Figure 4M**) Additionally, TAMs co-cultured with OVA specific CD8 cells, showed enhanced antigen presentation when cultured with a full OVA protein, which does not activate OTI CD8 T cells alone without an antigen presenting cell (**Extended Data Figure 10C**). Further, T cells within these tumors displayed reduced expression of coinhibitory markers PD-1, LAG3 and TIM3 expression (**Figure 4N-O, Extended Data Figure 10 D-E**). PD-1 is thus a direct suppressor of macrophage metabolism and function to promote anti-tumor immunity.

**Figure 4.**
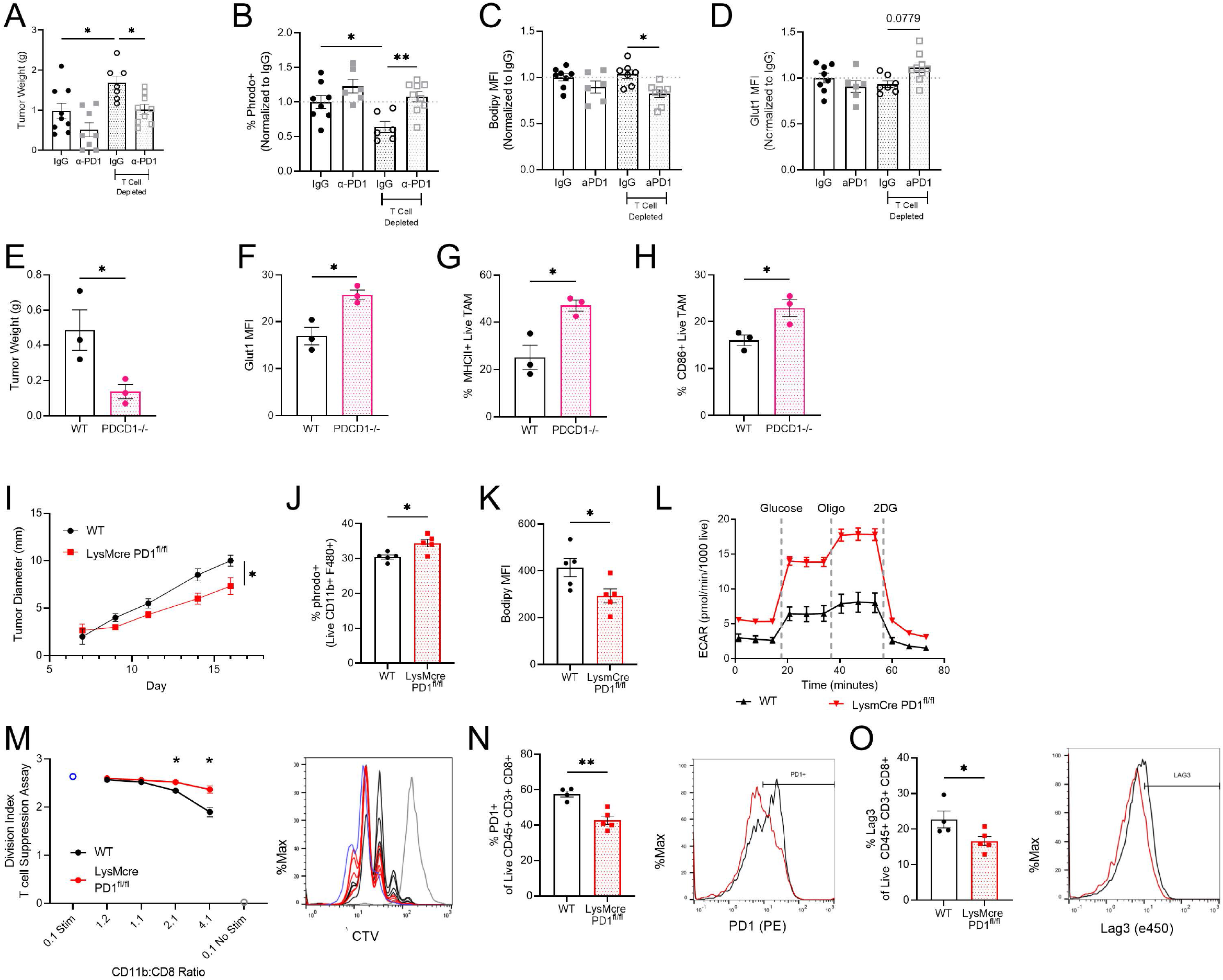
Targeting macrophage-specific PD-1 improves T-cell mediated anti-tumor immunity. A-D) Tumor weight (A) and CD11b^+^/F480^+^ phagocytosis (% phrodo+, B), fat ontent (Bodipy MFI, C), and GLUT1 expression (D) on day 16 in MC38 tumor in mice treated with anti-CD4 and anti-CD8 cocktail on days -1, 2, 7, 12, 15, and 200mg anti-PD-1 or IgG on days 9, 11, 13, 15. Data are combined from two independent experiments (n=6-9 mice) and B-D show data normalized to within-experiment IgG controls. E-H) Tumor weight (E) and GLUT1 (F), MHCII (G), and CD86 (H) expression on CD11b^+^ F480^+^ TAMs on day 14 following subcutaneous injection of 5×10^6^ MC38 cells injected into WT or *Pdcd1*-/-mice. Data representative of two independent experiments with 3-5mice per group. I) Tumor growth over time following injection of 10^6^ MC38 cells in WT vs LysMcre *Pdcd1*^fl/fl^ male mice. J-K) Percent of phrodo+ and bodipy MFI of TAMs within MC38 tumors on Day 16. L) GlycoStress Test of CD11b^+^ isolated cells from MC38 tumors. M) Division index calculated from flow cytometry analysis of CTV staining. CD11b+ isolated cells from MC38 tumors from WT or LysmCre *Pdcd1*^fl/fl^ mice were cocultured with WT CD8^+^ isolated splenocytes in the presence of 1mg SIINFEKL for 2 days. Representative histogram of CTV of CD8 cells. N-O) Percent of PD-1^+^ and Lag3^+^ CD8 T cells from MC38 tumors of WT and LysMcre *Pdcd1*^fl/fl^ mice with representative histograms. Data represent ≥ three independent experiments with 3-6 mice per group. Data are mean ± s.e.m. (ns p>0.05, *p≤0.05). P values were calculated using a one-way ANOVA (a-d) or an unpaired two-tailed t-test (e-o). ns p>0.05, *p≤0.05, **p≤0.001).

## The obesity-cancer connection is linked to PD-1 mediated macrophage dysfunction

Obesity both promotes tumor initiation and progression yet can also augment responses to immunotherapy, contributing to the observed paradox. These effects of obesity occur in part through direct action of cytokines and altered metabolites on cancer cells, but likely also occur through modulation of anti-tumor immunity. Our findings show that PD-1 itself contributes to this obesity-cancer connection as obesity-associated inflammatory cues upregulated PD-1 on macrophages. This was selective for TAMs, as PD-1 expression on tumor infiltrating T cells was not altered and splenic macrophages did not increase PD-1. An obesity-driven shift in immune cell populations has the potential to reshape the tumor immune microenvironment particularly through impaired macrophage function. DIO decreased the ability of macrophages to uptake and present antigen to T cells to ultimately reduce T cell-mediated immune surveillance that could allow more rapid tumor growth. Responses to the widely used PD-1 checkpoint therapy may subsequently be enhanced in obese individuals by dual action of releasing obesity-associated inhibition of macrophages together with increased T cell function when PD-1 is blocked. Importantly, PD-1 expression acts directly on macrophages to impact pro-inflammatory anti-tumor function and metabolism. These findings identify PD-1 as a metabolic regulator in TAM dysfunction and reveal a unique PD-1 mediated and macrophage-specific mechanism for immune tumor surveillance and checkpoint blockade. It is important to note that the direct effects of PD-1 were tested in settings where other innate immune cells, such as dendritic cells, may also be affected. Future work will be needed to explore the impacts on and by these other myeloid cells in the tumor microenvironment. In addition, it would be important to explore the role of combination therapies with this newly identified mechanism of anti-PD-1. It remains unclear whether these macrophage-mediated effects may provide prognostic indicators for immune checkpoint therapies. These findings provide mechanistic insight into obesity, high fat diet, and immune checkpoint therapies and the role of immune checkpoints in obesity associated cancers. Obesity associated inflammation and fatty acids provide a metabolic link in macrophage specific PD-1 expression. Taken together, we uncover an obesity-cancer connection through induction of PD-1 on macrophages and an alternative mechanism for the obesity paradox of immune checkpoint therapy, in which PD-1 blockade directly enhances TAM-mediated stimulation of anti-tumor T cells.

## Acknowledgements

We thank members of the J.C.R. and W.K.R. laboratories for their constructive input; the J. Balko, Y. Kim and P. Hurley laboratories for the use of their tumor dissociators; This work was supported by K00 CA234920 (J.E.B.), F30 CA239367 (M.Z.M.), F30 CA247202 (B.I.R.), R01 CA217987 (J.C.R.), NIH T32 GM007753 (B.T.D.), K08 CA241351 (S.M.H), R01 DK105550 (J.C.R.), the American Association for Cancer Research (B.I.R. and W.K.R.), T32 GM007347 (M.Z.M., B.I.R.), K12 CA090625 (K.E.B. and W.K.R.), and the Vanderbilt-Incyte Alliance (J.C.R. and W.K.R.). The Vanderbilt VANTAGE Core, including A. Jones, provided technical assistance for this work. VANTAGE is supported in part by a CTSA Grant (5UL1 RR024975-03), the Vanderbilt Ingram Cancer Center (P30 CA68485), the Vanderbilt Vision Center (P30 EY08126) and the NIH/NCRR (G20 RR030956). Flow-sorting experiments were performed in the VUMC Flow Cytometry Shared Resource by D. K. Flaherty and B. K. Matlock and were supported by the Vanderbilt Ingram Cancer Center (P30 CA68485) and the Vanderbilt Digestive Disease Research Center (P30 DK058404). We acknowledge the Translational Pathology Shared Resource supported by NCI/NIH Cancer Center Support Grant 5P30 CA68485-19 and the Vanderbilt Mouse Metabolic Phenotyping Center Grant 2 U24 DK059637-16. Figs. 1a were created with Biorender.com.

## Author contributions

J.E.B and J.C.R. conceived and designed the study and composed the manuscript. K.E.B. provided clinical expertise and human ccRCC samples for flow cytometry analysis. M.D.L assisted with collecting and processing of human patient samples. J.E.B., M.M.W., B.I.R., M.Z.M performed DIO and tumor experiments and assisted with downstream analysis including flow cytometry. G.A.N., M.A.C., X.Y analyzed scRNAseq data set. E.A.A. provided expertise in staining of tumor microarray samples that were collected from S.M.H.. E.S.H. and K.K.S assisted with experimental studies and editing the manuscript. V.A.B. generously provided transgenic LysMCre *Pcd1*^*fl/fl*^ breeding pairs. S.A.H. provided clinical expertise and provided tumor microarray samples. J.C.R. and W.K.R. obtained funding for this study.

