## Supplemental Figure Legends for "PD-1 is induced on tumor-associated macrophages in obesity to directly restrain anti-tumor immunity"

**Extended Data Figure Legends**

**Extended Data Figure 1. High Fat Diet Feeding induces an obese phenotype**

A) Mouse body weights during 20 weeks of LFD or HFD feeding prior to MC38 tumor injection (n=20 mice per diet). B) Mouse food intake during 20-week diet treatment. Food weighed weekly per cage and intake calculated as food weight difference between weeks (difference in food weight(g)/7 days/5 mice per cage X kcal of diet) (45% HFD=4.73kcal/g; LFD=3.85kcal/g) (n=4 cages with 5 mice per cage). C-D) Blood glucose and blood insulin levels measured via tail bleed following a 4-hour fast after 20 weeks of diet treatment (n=20 mice per diet). E-H) Representative tumor growth curves of mice inoculated with 5x10^5^ MC38 cells (n=6-8 mice per group). I) Tumor weights on day 19 following subcutaneous inoculation. “X” indicates days of anti-PD-1 or IgG injection. Data points identified as “+” were pooled together at equal ratios of live cells for subsequent single cell RNA sequencing analysis. J) Percent change in tumor growth in response to a-PD-1 treatment within respective diets ((Tumor Diameter Day 9 LFD a-PD-1 – Diameter Day 19 LFD IgG) ÷ Diameter Day 19 LFD IgG × 100). P values were calculated using an unpaired two-tailed t-test (A-D, J) or one-way ANOVA (I). ns=p>0.05; * p≤0.05; ** p≤0.01.

**Extended Data Figure 2. Cell characterization and metabolic signature of CD45^+^ and CD45^-^ cells from MC38 tumors**

A-B) Cell counts of SingleR annotated clusters from scRNAseq of CD45^-^ and CD45^+^ MC38 tumor samples. C) Heat map of identifying genes used to categorize CD45^+^ cells. D) Violin Plot of gene expression of *Cd44*, *Cd69*, *Icos*, and *Ifng* within CD8 identified clusters from CD45^+^ tumor fraction. E) UMAP of clone abundance within each treatment group from V(D)J Sequencing of T cell clusters with productive TCRs. F) Absolute number of T cells with paired TCR-a and TCR-b sequences. G) Volcano plot of Compass determined metabolic reactions within Fatty acid oxidation Recon2 pathway from CD8 and Macrophage identified clusters comparing LFD IgG to HFD IgG groups. Data is representative of one experiment, n = 3 pooled tumors from LFD mice and n = 3 pooled tumors from HFD mice. P values were calculated using wilcoxon rank-sum test (D) ns=p>0.05; * p≤0.05.

**Extended Data Figure 3. CD8 and TAM phenotypes in MC38 tumors following anti-PD-1 and HFD**

A) Representative gating strategy for quantifying T cell and macrophage cell populations within the spleen and tumor microenvironments. B) Quantification of tumor CD8^+^ cells gated on live singlets. C) Quantification of tumor PD-1^+^ CD8 T cells with representative histogram. Top gray histograms show splenic CD8 T cell PD-1 expression. D-E) Quantification of LAG3 and TIM3 expression on CD8 T cells. F) Quantification of F480^+^ cells gated on live singlets. G) Quantification of PD-1^+^ TAMs with representative histogram. H) Measurement of basal OCR, ECAR and OCR:ECAR ratio of CD11b^+^ sorted from MC38 tumors following a MitoStress Test. Each data point represents a biological replicate; data are mean ± s.e.m. Data represent ≥ three independent experiments with ≥ 3 mice per group. P values were calculated using a one-way ANOVA, ns=p>0.05; * p≤0.05.

**Extended Data Figure 4. Quantification of PD-1 expression in human TAMs**

A) Gating strategy of freshly isolated human PBMCs with matched ccRCC tumors pre-enriched for TILS and TAMs using CD8 and CD11b magnetic beads. B) Histogram of TIM3 and LAG3 expression in PBMC and tumor TAMs and CD8 cells. C) *PDCD1* expression within immune cells subsets from scRNAseq within TISCH2 database. D-E) *PDCD1* expression in CD8 T effector and Monocyte/Macrophage identified cells in the TISCH2 scRNAseq database from various cancer datasets. F) Percent *PDCD1*^+^ macrophage cells corresponding to pathological grade compared to adjacent normal tissue from human ccRCC tumor samples measured using gene expression from scRNAseq (n=11 patients). G-H) Percent *PDCD1*^+^ macrophages and percent within corresponding pathological grade of human CRC tumor samples quantified using gene expression from scRNAseq (n=62 patients). I) Enrichment of KEGG metabolic signature scores of *PDCD1*^-^ versus *PDCD1*^+^ TAMs from human CRC scRNAseq. C-D each data point represents a separate single cell data set from congregated database within TISHC2; F-H each data point represents individual patient samples sequenced. Data are mean ± s.e.m. P values were calculated using an unpaired two-tailed t-test (G) or a one-way ANOVA (H). ns p>0.05, **p≤0.01.

**Extended Data Figure 5. Characterization of PD-1 expression in mouse macrophages**

A-B) Normalized transcript counts of *Pdcd1* and *Cd274* in the indicated MACS-sorted cell populations in MC38 tumor samples. C) Percent of *Pdcd1*^+^ macrophages within MC38 tumor scRNAseq transcriptomes. D) Percent of CD11b^+^ F480^+^ macrophages within MC38 tumors and spleens following 2 weeks of LFD or HFD treatment (acute diet). E) Percent of PD-1^+^ macrophages in tumor and spleen following 2-week acute diet treatment. F) Gating strategy tumor immune cells. G-H) Relative abundance of indicated immune cell populations in tumor (G) and spleen (H) with diet and aCD4/8 treatments (n=3 mice/group) with representative flow cytometry plots demonstrating T cell depletion. I) CD274 (PD-L1) expression of BMDMs following 4-, 8-, 24- and 48-hour exposure to IFNg (M1), IL-4 (M2) or tumor-conditioned media (TCM). J-K) Mean fluorescence intensity of PD-1 (J) and PD-L1 (K) expression and representative histograms of BMDMs following 24-hour exposure to LPS, IFNg, IL-4, TCM, IL-6, MCP-1, IL-1b, TNF-a, leptin, or insulin. Data in A-B is from one independent flow-sort Nanostring experiments. Data in C is representative of one experiment, n=3 pooled tumors from LFD or HFD mice. Data in H is of 3-5 pooled spleens per treatment group. Data points in G, I-K indicate biological replicates (n=3-5). Data are mean ± s.e.m. P values were calculated using an unpaired two-tailed t-test (d-e) or a one-way ANOVA (g, i-k). ns p>0.05, *p≤0.05).

**Extended Data Figure 6. PD-1+ TAMs have a unique gene signature**

Differential gene expression of select genes from *Pdcd1*^-^ versus *Pdcd1*^+^ TAMs from MC38 scRNAseq. See Extended Data Table 1 for full list of significant genes

**Extended Data Figure 7. Flow cytometry gating strategy for PD-1 expression on macrophages**

A) Representative gating strategy of FACS sorted PD-1 hi and PD-1 lo TCM-treated BMDMs. B) Gating strategy of PD-1 hi and PD-1 lo CD11b^+^/F480^+^ macrophages from MC38 tumors.

**Extended Data Figure 8. Direct effects of anti-PD-1 treatment on macrophage metabolism**

A) MFI of 2-NBDG in peritoneal macrophages following 45min exposure to 2-NDBG (n=9 mice). B-C) OCR and ECAR from MitoStress Test of BMDM from WT or *Pdcd1*-/- mice stimulated for 24 hours with IFN-g (M1), IL-4 (M2), or TCM. Data points represent biological replicates (n=6mice) D-F) PD-1 expression of stimulated bm macrophages from WT mice treated with 2DG (5mM), DON (5mM) or etomoxir (100mM) for 12 or 24 hours (n=3 mice) G-H) OCR and ECAR from MitoStress Test of BMDM stimulated with TCM for 24 hours followed by treatment with rmPD-L1 and/or anti-PD-1 for an additional 24 hours. I-J) Baseline ECAR and maximal ECAR measured from GlycoStress test of indicated treated bm macrophages. Data represent ≥ three independent experiments with 5 mice per group. Data are mean ± s.e.m. (ns p>0.05, *p≤0.05). P values were calculated using an unpaired two-tailed t-test (b) or a one-way ANOVA (e-g, j-k). ns p>0.05, *p≤0.05).

**Extended Data Figure 9. Antibody treatment effectively depletes CD4 and CD8 T cells in the spleen and tumor microenvironment.**

A) Representative gating strategy for tumor immune cell subsets. B) Tumor immune cell populations following anti-CD4/CD8 depletion and/or anti-PD-1 treatment from MC38 tumors on Day 16. C) Representative flow cytometry data of T cell depletion in tumor and spleen of anti-CD4/CD8 treated mice. Data point represents biological replicates (n=3-4) mice per group. Data are mean ± s.e.m. P values were calculated using a one-way ANOVA (p>0.05, *p≤0.05, **p≤0.001).

**Extended Data Figure 10. Myeloid specific PD-1 depletion improves TAM anti-tumor function**

A) Representative gating strategy for tumor immune cell subsets. B) Percent of immune cell subsets from MC38 tumors injected into WT and LysMcre *Pdcd1*^fl/fl^ mice. C) Division index calculated from flow cytometry analysis of CTV staining. CD11b^+^ isolated cells from MC38 tumors from WT vs LysmCre *Pdcd1*^fl/fl^ mice cocultured with WT CD8^+^ isolated splenocytes. Cocultured cells treated in the presence of 1mg/ml OVA protein for 5 days. D-E) Percent and MFI of TIM3 and CD69 expression on CD8 T cells from MC38 tumors of WT and LysmCre *Pdcd1*^fl/fl^. F) MitoStress Test of CD11b^+^ isolated cells from MC38 tumors of WT and LysmCre *Pdcd1*^fl/fl^ with basal and maximal OCR measurements. G) Calculated glycolysis and glycolytic capacity following GlycoStress Test of CD11b^+^ isolated cells from MC38 tumors of WT and LysmCre *Pdcd1*^fl/fl^. Data represent ≥ three independent experiments with 3-6 mice per group. Data are mean ± s.e.m. P values were calculated using an unpaired two-tailed t-test. (ns p>0.05, *p≤0.05, p≤0.01).
