## Supplementary material for "PD-1 is induced on tumor-associated macrophages in obesity to directly restrain anti-tumor immunity": Methods

**Mice**

C57BL/6J (Stock #000664), *Pdcd1^-/-^* (Stock #028276), and OTI transgenic (Stock #003831) mice were purchased from The Jackson Laboratory. *LysMcre Pdcd1^fl/fl^* mice were kindly provided by Dr. Vassiliki A. Boussiotis of Harvard Medical School. All mouse procedures were performed under Institutional Animal Care and Use Committee (IACUC)-approved protocols from VUMC and conformed to all relevant regulatory standards. Mice were housed in ventilated cages with at most five mice per cage and were provided with ad libitum food and water. Mice were on 12-h light–dark cycles which coincided with daylight in Nashville, TN. The mouse housing facility was maintained at 20–25°C and 30–70% humidity.

**Diet Treatment**

For DIO experiments, mice starting at 6-10 weeks old were fed a 45 kcal% fat diet (Research Diets D12451) or a 10 kcal% sucrose matched control diet (Research Diets, D12450H). Mice were maintained on their respective diet for 20-25 weeks before initiating injectable tumor studies. Metabolic parameters were assessed at 19-21 weeks of dietary treatment. After a five-hour fast, blood samples were collected from the tip of the tail. Blood glucose concentrations were determined in whole blood using a glucometer (Bayer Contour, Michawaka, IN). Collected blood was centrifuged at 4,000 rpm for 10 min at 4°C.  Plasma was aliquoted and stored at -80°C until analysis. Plasma insulin (Mercodia Inc., Winston Salem, NC) concentrations were analyzed using commercial ELISA kit according to the manufacturer’s instructions. For injectable tumor models, mice were injected subcutaneously in the posterior flank with 5x10^5^ MC38 cells. For non-DIO experiments, mice were injected subcutaneously with 2x10^6^ MC38 cells. Once palpable tumors were present, tumor measurements were performed using a caliper every 2-3 days. Mice were euthanized if the IACUC-approved humane end-points were reached. In immunotherapy studies, tumor-bearing mice received intraperitoneal injections of either 200 μg anti-mouse PD-1 Ab (RMP1-14; BioXCell) in 200 μl PBS or 200 μg rat IgG2a isotype control (2A3; BioXCell) in 200 μl PBS on days 9, 11, 13, and 15 post tumor injection. For antibody-mediated T cell depletion, mice were treated with depleting antibodies or isotype control delivered by intraperitoneal injection on days −1 (200 μg), 2 (200 μg), 7 (200 μg), 12 (200 μg), and 16 (200 μg) relative to tumor injection (day 0). For T cell depletion experiments, the following antibodies were used: rat IgG2b isotype control (BioXCell, Clone 2A3), anti-CD8α (BioXCell, Clone 2.43), and anti-CD4 (BioXCell, Clone GK1.5)

**Cell lines and primary macrophage isolation**

MC38 cells were grown in RPMI supplemented with 10% FBS and 1% pen/strep at 37°C in a humidified 5% CO_2_ incubator. All FBS was heat-inactivated prior to use. For primary macrophage experiments, bone marrow was flushed from femurs of 6-10 week-old mice and incubated in the presence of 20ng/mL M-CSF (Peprotech) for 5 days. Bone marrow derived macrophages (BMDMs) were stimulated with 50ng/mL IFN-g, IL-4, or 25% MC38-conditioned media for 24 hours before further analysis. Following stimulation, BMDMs were treated with 1μg anti-mouse PD-1 Ab (RMP1-14; BioXCell) followed by 1μg recombinant mouse PDL1/B7-H1 His Tag Protein (R&D Systems, 9048-B7) for an additional 24 hours before flow cytometry analysis or metabolic flux assay. For metabolic inhibitor assays, macrophages were treated following stimulation for an additional 24 hours with DON (5μM, Sigma-Aldrich D2131), 2-deoxyglucose (2DG, 5mM, Cayman Chemical 14325), or etomoxir (100μM, Cayman Chemical, 11969) or equal volume PBS vehicle were added. Conditioned media was obtained by growing MC38 cells to 90% confluence and treating with 50ng/mL IFNγ for 24hours. Following treatment, cells were washed and incubated with fresh media for another 24 hours. Supernatant of treated MC38 cells were sterile filtered through 0.45uM filter and frozen into small aliquots for subsequent use. Peritoneal macrophages (pMacs) were isolated by flushing peritoneal cavity of 6-10week old mice with 10mL of media. Peritoneal exude cells were cultured in tissue culture treated plates in MCSF containing media for 24hours. Non-adherent cells were washed away following 24hours, and adherent cells were identified as peritoneal macrophages and used in subsequent analysis. To remove adherent BMDM or pMacs, cells were washed with PBS and incubated in 0.05% Trypsin for 5min before being gently scraped with a cell scraper.

**Tissue dissociation**

Mice were euthanized and spleen and tumors were collected as previously described^37^^,^^38^. Single-cell suspensions of splenocytes were prepared by mechanical dissociation followed by ACK-lysis. Tumors were chopped, mechanically dissociated on the Miltenyi gentleMACS Octo Dissociator with Heaters (setting implant tumor one) and digested in 435 U ml^−1^ DNase I (Sigma-Aldrich, D5025) and 218U ml^−1^ collagenase (Sigma-Aldrich, C2674) at 37 °C for 30 min. After enzyme treatment, tumors were passed through a 70-μm filter and ACK-lysed. Cells were resuspended in complete RPMI and counted using trypan blue with the TC20 Automated Cell Counter (Bio-Rad). Next, cell suspensions were fractionated using magnetic bead positive selection according to the manufacturer’s instructions (all Miltenyi mouse kits: CD45 TIL 130-110-618, Dead cell removal kit 130-090-101, CD11B 130-049-601). In brief, cells were resuspended at 10 million total cells per 90 μl MACS buffer and 10 μl microbeads for 15 min. Then, cell suspensions were applied to LS columns (Miltenyi, 130-042-401) in Miltenyi QuadroMACS Separators, washed and eluted according to the manufacturer’s instructions.

**Mouse flow cytometry**

Single-cell suspensions obtained from tumors, spleens or primary cell culture were incubated in F_c_ block (1:50, BD, 553142) for 10 min at room temperature, stained for surface markers for 15 min at room temperature, washed with FACS buffer (PBS + 2% FBS) once and resuspended in FACS buffer for analysis on a Miltenyi MACSQuant Analyzer 10 or 16. For intracellular staining, the eBioscience FOXP3/transcription factor staining buffer kit (Thermo Fisher Scientific, 00-5523-00) was used. Surface staining was performed as described above, cells were fix/permed for 20 min at 4 °C, and then stained for intracellular markers for at least 30 min at 4 °C. Ghost Dye Red 780 viability dye (1:4,000, Cell Signaling, 18452S) was used identically to surface antibodies. Anti-mouse and cross-reactive antibodies used were: CD45 BV510 (1:1,600, 30-F11, Biolegend, 103138), B220 e450 (1:400, RA3-6B2, Thermo Fisher Scientific, 48-0452-82), CD11B e450 (1:1,600, M1/70, Thermo Fisher Scientific, 48-0112-82), CD11B FITC (1:1,600, M1/70, Biolegend, 101206), CD8A AF488 (1:1,600, 53-6.7, Biolegend, 100723), CD8A BV510 (1:600, 53-6.7, BD, 563068), LY6C FITC (1:4,000, HK1.4, Biolegend, 128006), CD11C PE (1:1,000, N418, BioLegend, 117308), FOXP3 PE (1:125, FJK-16 s, Thermo Fisher Scientific, 12-5773-82), CD4 PerCP-Cy5.5 (1:600, RM4-5, BioLegend, 100540), LY6G PerCP-Cy5.5 (1:800, 1A8, BioLegend, 127616), F4/80 PE-Cy7 (1:800, BM8, BioLegend, 123114), NKp46 PE-Cy7 (1:200, 29A1.4, BioLegend, 137618), CD3 PE-Cy7 (1:200, 17A2, BioLegend, 100220), CD3 FITC (1:200, 17A2, BioLegend, 100204), CD3 APC (1:200 17A2, BioLegend, 100236), CD206 APC (1:500, C068C2, BioLegend, 141708), GLUT1 AF647 (1:500, EPR3915, Abcam, ab195020), MHCII I-A/I-E APC (1:4,000, M5/114.15.2, BioLegend, 107614), LAG3 e450 (1:100, eBioC9B7W, Thermo Fisher Scientific, 48-2231-82), PD-1 PE (1:100, 29F-1A12, BioLegend, 135206), TIM3 APC (1:100, RMT3-23, BioLegend, 119706), CD44 PE-Cy7 (1:1,000, IM7, BioLegend, 103030). CD80 (1:200, 16-10A1, BioLegend, 104713), CD86 APC (1:200, GL-1, BioLegend, 10511), MHCII FITC (1:400, BioLegend, 107605), PDL1 BV605 (1:400, BioLegend, 124321), MHCI FITC (1:800, BioLegend, 107605). The anti-human antibodies used were: CD45 BV421 (1:400, HI30, BioLegend, 304032), CD3 APC (1:200, UCHT1, BioLegend, 300439), CD11B PerCP-Cy5.5 (1:200, ICRF44, BioLegend, 301328), CD14 BV510 (1:200, M5E2, BioLegend, 301842), and Human Fc Block (1:50, BD 564220). For *ex vivo* fluorescent palmitate or glucose uptake, tumor single-cell suspensions were acclimated for 45 minutes in XF RPMI with 200mM L-Glutamine (Agilent 103681-100) at 37 °C, 5% CO_2_, incubated with BODIPY FL C16 (1μM in XF RPMI), 2NDBG (100μM Sigma A8625), for 45 min, washed twice with FACS and then stained for surface markers. For phagocytosis assay, single-cell suspensions were incubated for 45min with 100μg/ml of phrodo red *E coli* beads (Invitrogen P35361)

For myeloid suppression assays, microbead-isolated CD11B^+^ myeloid cells were plated in 96 well plates and serially diluted starting at 4x10^5^ cells/well. Microbead-isolated CD8^+^ splenocytes (1x10^5^ cells/well) from OTI TCR-transgenic mice were isolated using a negative isolation kit (Miltenyi 130-095-236) and were stained with CellTrace Violet and plated in the CD11b plated wells at 10^5^ cells/well in addition to 1 μg/ml SIINFEKL peptide (Sigma-Aldrich S7951). CellTrace Violet (CTV) (Thermo Fisher Scientific, C34557) was used at 1:1000 for cell proliferation assays. CD11b/CD8 co-cultures were incubated in DMEM containing 5% FCS, 2 mM glutamine, 100 units/mL penicillin-streptomycin, 10 mM Hepes and 20 μM beta mercaptoethanol for 48-72 hr. As positive and negative controls OTI CD8^+^ splenocytes alone were incubated with 1 μg/ml SIINFEKL peptide or no peptide, respectively. Flow cytometry data were analyzed using FlowJo v.10.7.1. For antigen presentation assay, CD11b/CD8 co-cultures were prepared identically to suppression assay and incubated with 1mg/mL OVA protein (Ovalbumin, 323-329, Japanese quail, Sigma, O1641) for 5 days before analysis via flow cytometry.

**Extracellular flux assay**

For isolated TAMs, BMDMs and peritoneal macrophages, cells were plated at 50,000-100,000 live cells per well in at least 4 technical replicates on a Cell-Tak-coated plate (Corning 354240) in Agilent Seahorse RPMI 1640 supplemented with 10 mM glucose, 1 mM sodium pyruvate and 2 mM glutamine (for GlycoStress test RPMI without glucose and sodium pyruvate was used). Cells were analyzed on a Seahorse XFe 96 bioanalyzer using the MitoStress assay with 1 μM oligomycin, 2 μM FCCP and 0.5 μM rotenone and antimycin A or the GlycoStress assay with 10mM glucose, 1μM oligomycing, and 50mM 2-DG. Data were analyzed in Agilent Wave software v.2.6. For cell normalization, following MitoStress or GlycoStress assays, cells were stained with propidium iodide (eBioscience 00-6990-50) and Hoechst (ThermoFischer 62249) for 15min. Dead cells and total cells were counted using Cytation 5 and live cell number was calculated as total (Hoechst positive) – Dead (PI positive) cells before being imported into the wave normalization software and OCR/ECAR data presented as per 1000 live cells.

**TMA staining**

Human ccRCC TMA were provided by Scott Haake and Vanderbilt University Medical Center. Paraffin-embedded TMA slides were prepared for immunofluorescence and stained with anti-PD-1 (abcam #52587; 1:250), anti-CD163 (abcam #189915; 1:1,000), and anti-CD8 (Cell Signaling #70306S; 1:500) as previously described^39^. Briefly, slides were deparaffined in xylene and rehydrated in serial ethanol dilutions. Antigen retrieval was performed by heating slides for 17 min in Tris EDTA buffer, pH 9 in a pressure cooker at 110˚C. Slides were cooled to room temperature and then blocked with 2.5% horse serum (vector labs). After blocking, slides were incubated overnight at 4˚C with primary antibody in horse serum. Slides were then incubated in anti-rabbit or mouse HRP secondary (vector labs) for 1hr at room temperature the following day and subsequently incubated in 1:500 Opal 520, Opal 570, or Opal 590 (Akoya) for 10 minutes. For serial staining, slides were stripped using Citric Acid buffer, pH 6.1 in a pressure cooker at 110˚C for 2min and then staining was repeated using different antibody and Opal fluorophore. After last Opal staining, slides were mounted using antifade gold mount with DAPI (Invitrogen). Stained images were acquired using an Aperio Versa 200 Automated Slide imaging system (Leica/Aperio) via the Vanderbilt University Medical Center Digital Histology Shared Resource (DHSR) core. Images were analyzed with Fiji software. Quantification of markers was done by measuring total amount of fluorescence divided by total area of tissue (determined by H&E stain).

**Patient samples**

Fresh histology-confirmed ccRCC tumors and matched healthy tissue were surgically removed. Tumors and matched healthy kidney tissue were processed by mechanical dissociation (human tumor setting two on a Miltenyi gentleMACS) in Hanks’ balanced salt solution (HBSS) with calcium chloride and magnesium chloride. Mechanical dissociation was followed by enzymatic digestion in 435 U ml^−1^ DNase I (Sigma-Aldrich, D5025) and 218 U ml^−1^ collagenase (Sigma-Aldrich, C2674) in RPMI supplemented with 10% FBS, 1% glutamine, 1% penicillin–streptomycin, 1% HEPES and 0.1% 2-mercaptoethanol for 30–45 min, depending on tissue toughness, at room temperature with agitation at 17 rpm. Tissue digests were washed with HBSS without calcium chloride, magnesium chloride or magnesium sulfate and then incubated in 5 mM EDTA for 20 min at room temperature with agitation at 17 rpm. Tumor and matched healthy kidney digests were washed with HBSS with calcium chloride and magnesium chloride. Then they were passed through a 70-μm filter and ACK-lysed. Patient PBMCs were isolated by density gradient centrifugation using Ficoll-Paque (GE Healthcare, 17144002) in SepMate-50 tubes (StemCell Technologies, 85450) and subsequently ACK-lysed. Single-cell suspensions were frozen in 90% FBS, 10% DMSO. Batched tumor and matched PBMCs were thawed, rested for 10 min at 37 °C, counted, stained, and analyzed for flow cytometry. All studies were conducted in accordance with the Declaration of Helsinki principles under a protocol approved by the Vanderbilt University Medical Center (VUMC) Institutional Review Board (protocol no. 151549). Informed consent was received from all patients before inclusion in the study by the Cooperative Human Tissue Network at VUMC.

**Single Cell RNA Sequencing (scRNAseq)**

Eight-week-old C57BL/6 male mice were fed ad libitum with either a 10% kcal from fat (low-fat) diet or a 45% kcal fat (high-fat) diet (Research Diets D12450H and D12451, respectively). After 28 wk of diet treatment, mice were injected s.c. with 2 × 10^5^ murine MC38 (Kerafast) cells. Tumors were collected on day 20 post injection and then mechanically and enzymatically digested using mouse tumor digestion kit (Miltenyi) following the manufacturer’s instructions. Tumor-infiltrating leukocytes were sorted by positive selection using CD45^+^ microbeads (Miltenyi Biotec) from dissociated tumors for single-cell analysis. CD45^+^ cells were further enriched for live cells using a dead cell removal kit (Miltenyi Biotec). Cells were diluted with trypan blue and counted using a hemocytometer. Three tumors possessing the median weights were pooled together for each treatment group. Pooled samples were resuspended at 1 × 10^6^ cells/ml in PBS plus 0.4% BSA with a target of 20,000 live cells loaded onto the Chromium Controller (10x Genomics) and processed according to the manufacturer’s instructions. Sequencing was performed on the Illumina NovaSeq 6000 targeting 50,000 reads per cell for the 5' assay. The raw data (FASTQ files) were processed using the velocyto (version 0.17.17) to generate gene expression matrix as loom files.

Data preprocessing, normalization, integration, and clustering were performed within scanpy (version 1.7.1). In brief, the gene expression matrix for each sample was filtered by keeping cells with more than 200 but less than 4000 detected genes, as well as genes that were detected in more than three cells. Cells with mitochondrial genes representing greater than 10% of the transcripts were also removed. The matrix for each sample was integrated using a mutual nearest neighbor algorithm with 4002 highly variable genes (highly variable genes in at least two samples). Clustering was performed via Leiden on the integrated matrix, which identified 20 clusters that were then annotated using SingleR and manually confirmed. Differential gene expression analysis was done in scanpy using function “scanpy.tl.rank_genes_groups” with Wilcoxon rank-sum test to get significance. Violin plots of genes were made with scanpy. Data are available online at Gene Expression Omnibus (GSE179936).

For PD-1 analysis in human macrophages, a previously published dataset sequenced using the 10X Genomics platform from Zhang et. al. was obtained and converted to a Seurat v4 object^40^^,^^41^. Macrophages were then subset and labeled as PD-1^+^ or PD-1^-^ based on normalized expression of *PDCD1*. For differential expression between PD-1^+^ and PD-1^-^ macrophages, the FindMarkers function from Seurat v4 with the MAST test was used^42^. Significant differentially expressed genes were identified by an adjusted p value < 0.05.

**V(D)J sequencing**

Raw V(D)J sequencing files were processed using CellRanger (10X Genomics). Consensus V, D, and J gene sequences (i.e contigs) generated by CellRanger were further analyzed in R v4.2.1 by selecting for productive TCRs. In cases where multiple TCR were observed for a given cell, the most enriched TCR was retained. TCR results were incorporated into Seurat v4 objects generated using the cell by gene matrix from scanpy. Clones were called based on unique paired TCR-α and TCR-β sequences.

**Downstream analysis of Compass scores**

Single Cell RNA sequencing data as described above was further analyzed using Compass to infer the metabolic status of cells based on transcriptome data and flux balance analysis^27^. Compass was run - without micro pooling - four separate times with four separate inputs: LFD CD8, HFD CD8, LFD macrophage, and HFD macrophage using clusters classified from single R as macrophage or CD8. Compass and postprocessing of samples were followed as described in Wagner et al. Data presented as the -log of the reaction penalties, which makes the higher scores correspond with more active reactions and any reactions that were close to constant were removed (<10^-3^). Significant reaction values were determined using an unpaired Wilcoxon rank-sum test between the lean and obese mice of each group (CD8 and Macrophage) and all non-core reactions were filtered out. A meta-reaction was defined as belonging to core metabolism if it contains at least one core reaction. Core pathways are defined as Recon2 subsystems that have at least 3 core reactions. Metabolic genes are defined as the set of genes annotated in Recon2. Compass uses the package “Recon2”, which ranks confidence on a scale of 1-4 with 4 being the most confident, while a value of 0 means the confidence was not evaluated, only reactions with a confidence value of 4 or 0 were included, in addition all reactions in the citric acid cycle subsystem that occur outside of the mitochondria were removed. To decide which Recon2 subsystems were plotted, we first filtered both groups for subsystems having > 5 reactions. We further filtered the cd8 group by taking the median adjusted p-value of each subsystem and only keeping reactions whose medians were below the median of the medians (taking the most significant half of the subsystems) as well as the “Fatty acid oxidation” and “Fatty acid synthesis” subsystems. For the dot plots, we only plotted reactions shared between this filtered cd8 group and the unfiltered macrophage group whose subsystem had a median adjusted p-value of below 0.1. The color of the dot corresponds to the sign of the cohen’s d from the wilcoxin rank-sum test (blue = neg, red = pos), while the opacity of the dots correspond to their adjusted p-values (adjusted p    0.1 = faded).

**Nanostring Transcript Analysis**

Tumor cell suspensions were fractionated using serial magnetic bead positive selection according to the manufacturer’s instructions (all Miltenyi mouse kits: CD45 TIL 130-110-618, CD4/CD8 TIL 130-116-480, CD11B 130-049-601, F4/80 130-110-443). In brief, cells were resuspended at 10 million total cells per 90 μl MACS buffer and 10 μl microbeads for 15 min. Then, cell suspensions were applied to LS columns (Miltenyi, 130-042-401) in Miltenyi QuadroMACS Separators, washed and eluted according to the manufacturer’s instructions. RNA was isolated from tumor immune cell populations and unstained whole-tumour single-cell suspensions using the Quick-RNA Microprep Kit (Zymo R1050) according to the manufacturer’s instructions. RNA transcripts were quantified using the NanoString nCounter Metabolic Pathways Gene Expression Panel (XT-CSO-MMP1-12) according to the manufacturer’s instructions as previously described^38^.

**Statistics**

Prism software (GraphPad Software) was used to create graphs and conduct statistical analyses. Data were expressed as mean ± SEM. Analyses of the differences between two test groups were performed using the non-parametric unpaired two-way T test. For analysis of three or more groups, one-way ANOVA tests were performed with Tukey post hoc test. For scRNAseq violin plot, a Student *t* test was performed and adjusted using Benjamini–Hochberg procedure. The *p* values were considered statistically significant if *p* < 0.05.
