## Supplementary figures and images for "PD-1 is induced on tumor-associated macrophages in obesity to directly restrain anti-tumor immunity"

### Supplemental Figures

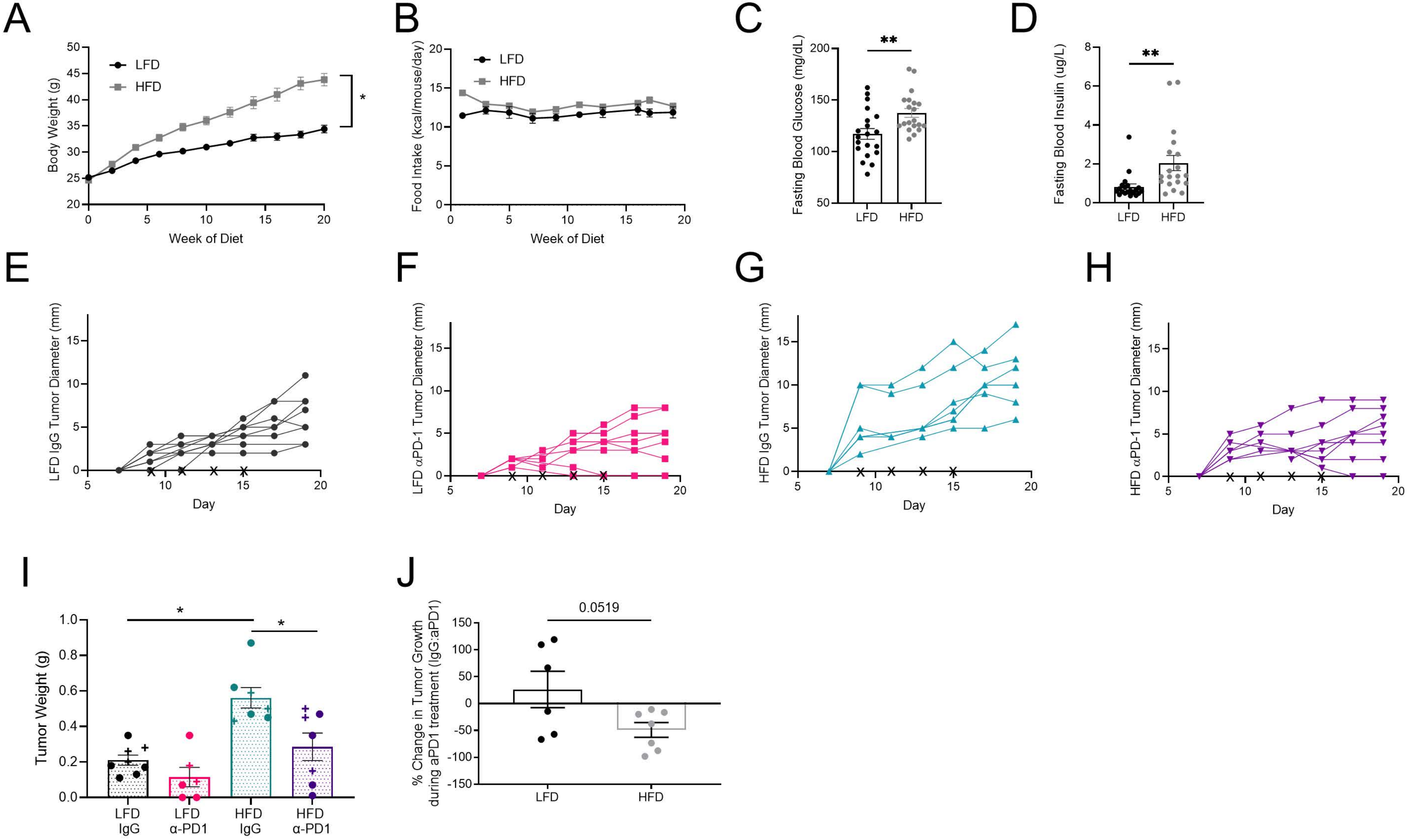

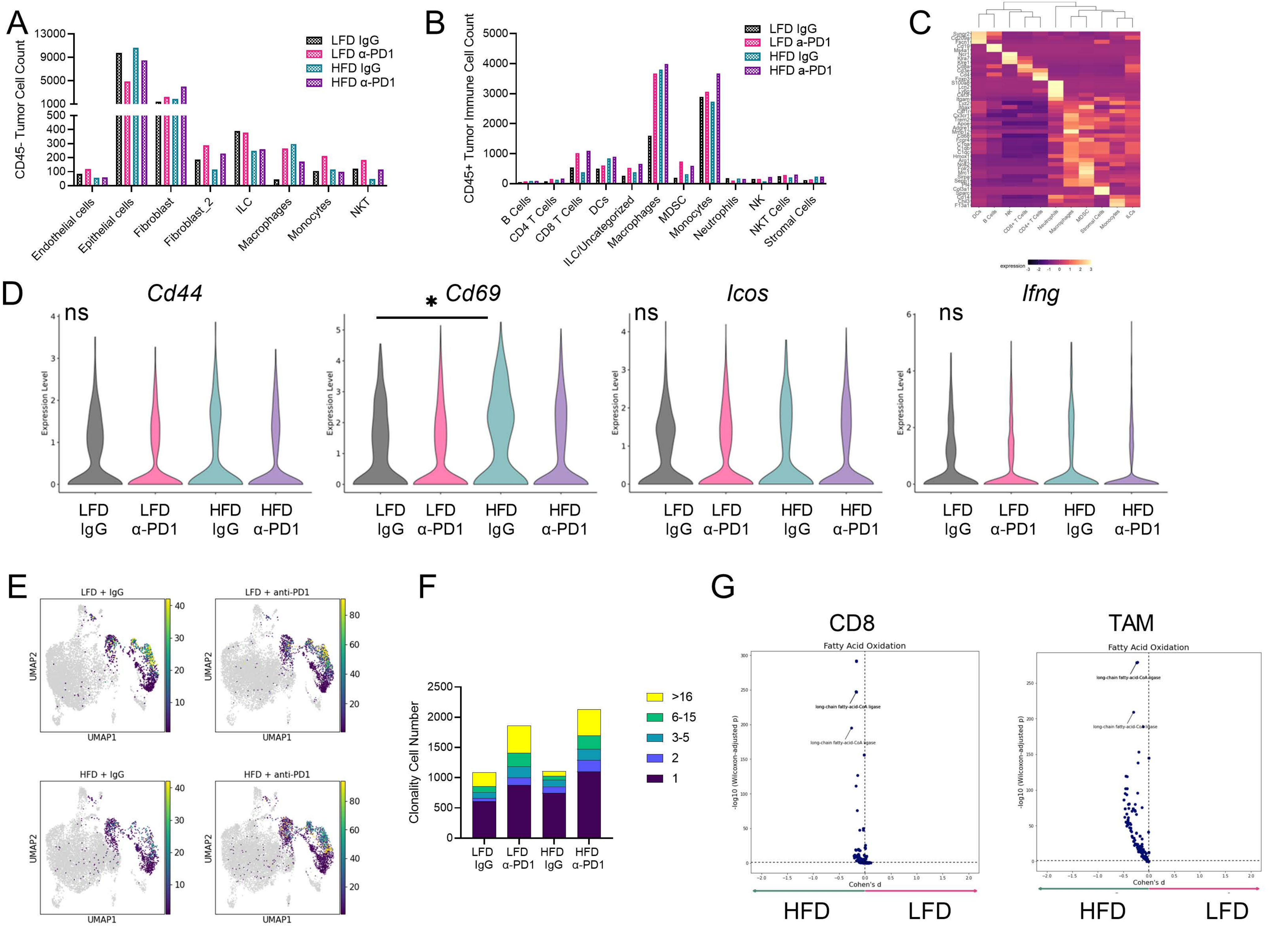

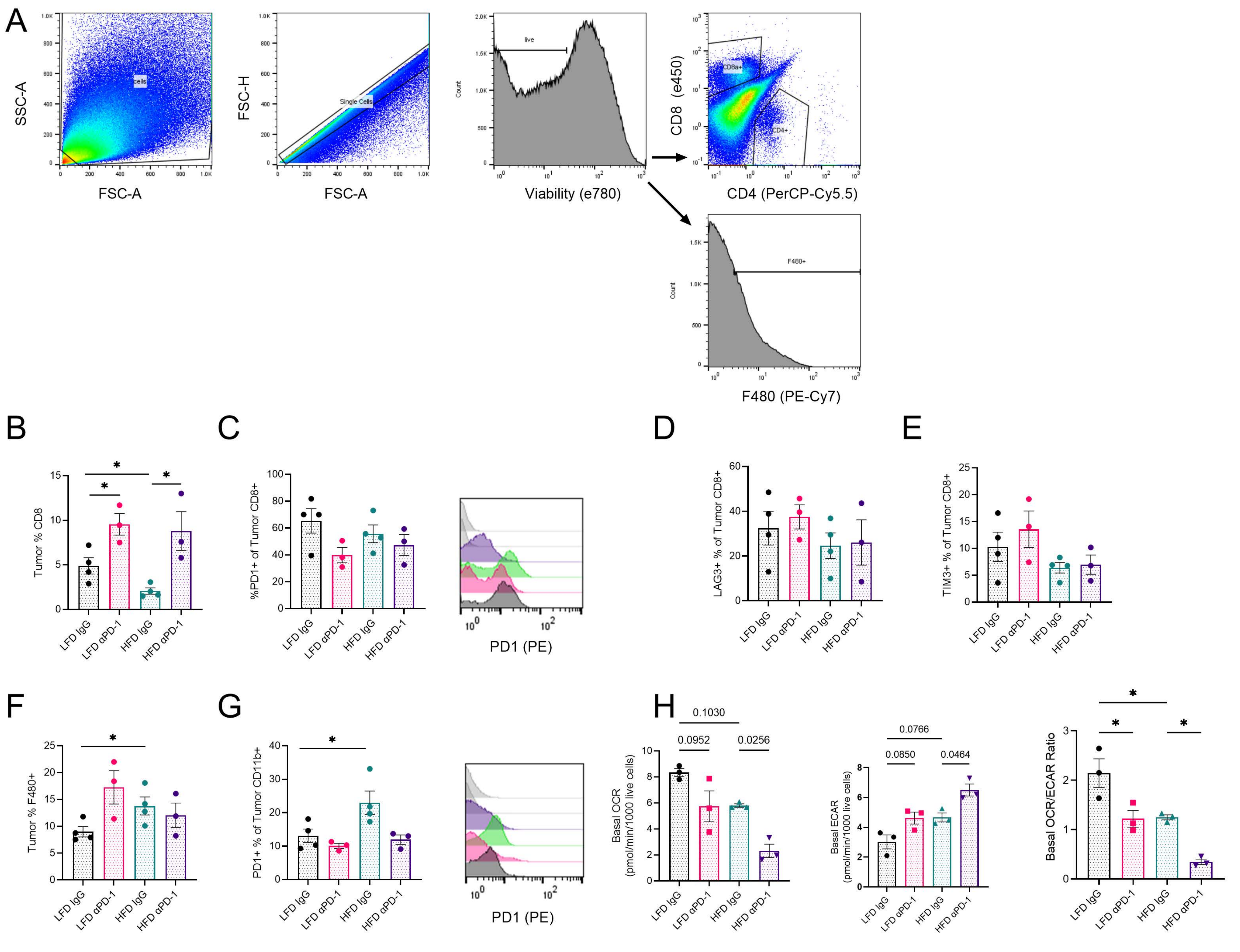

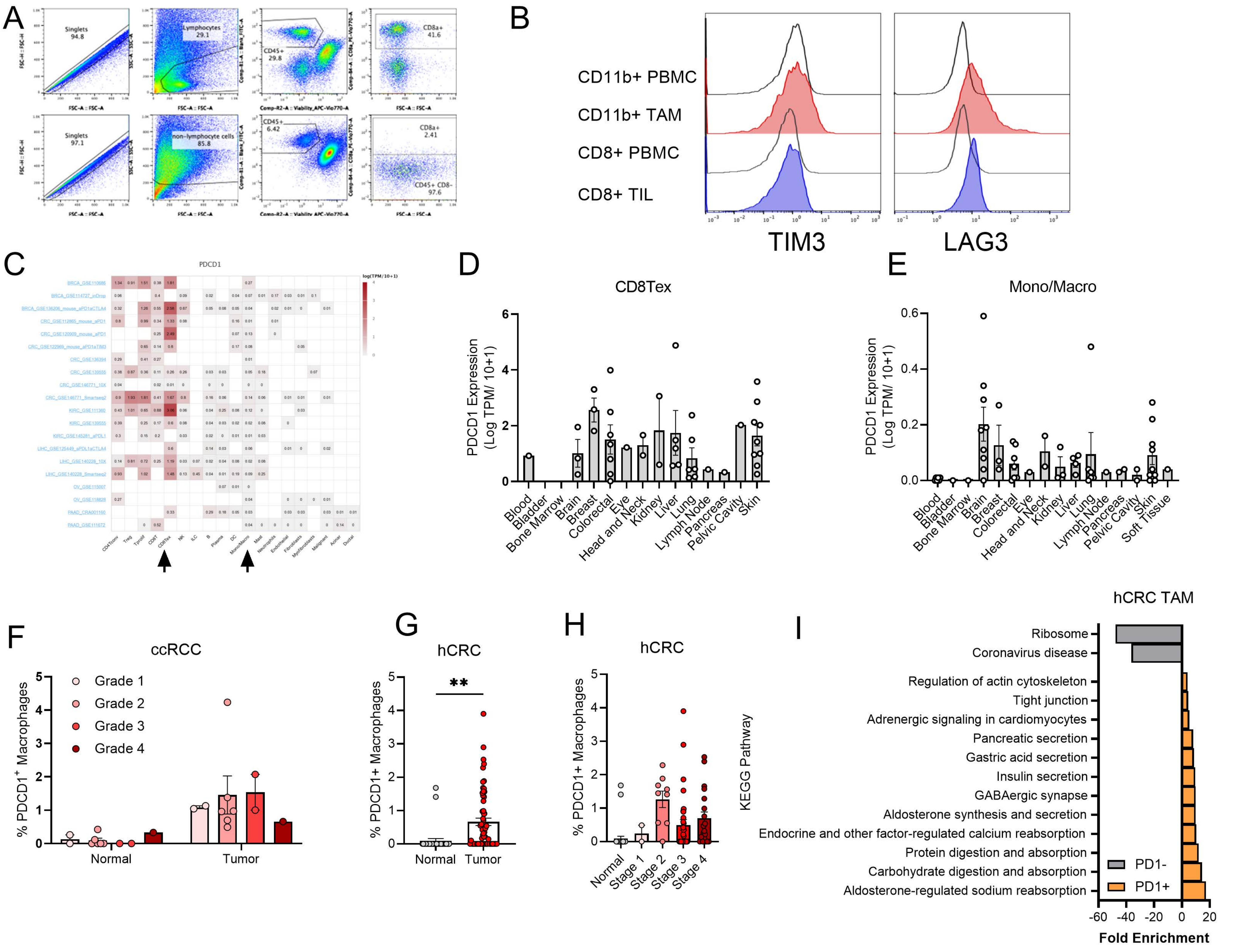

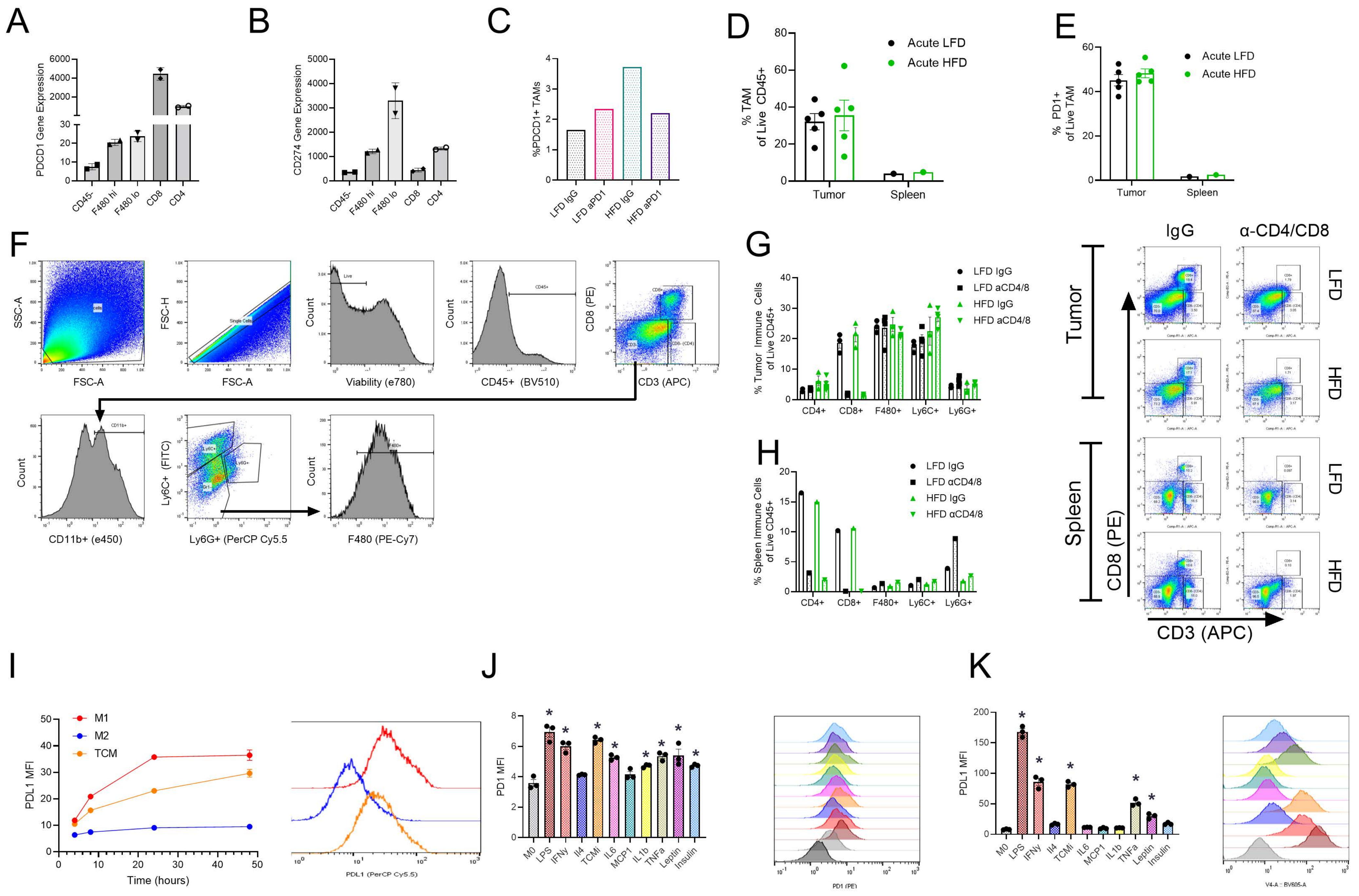

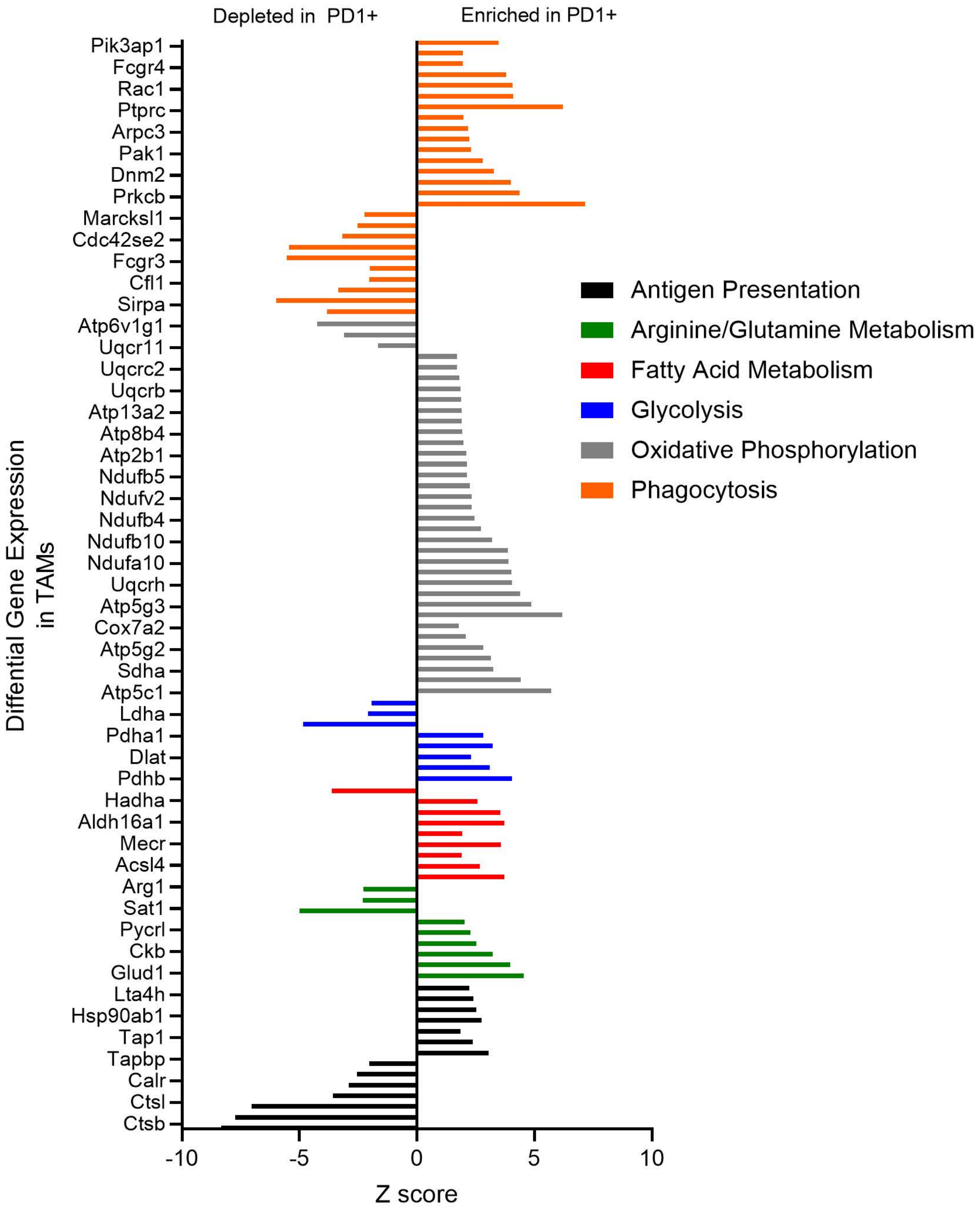

A

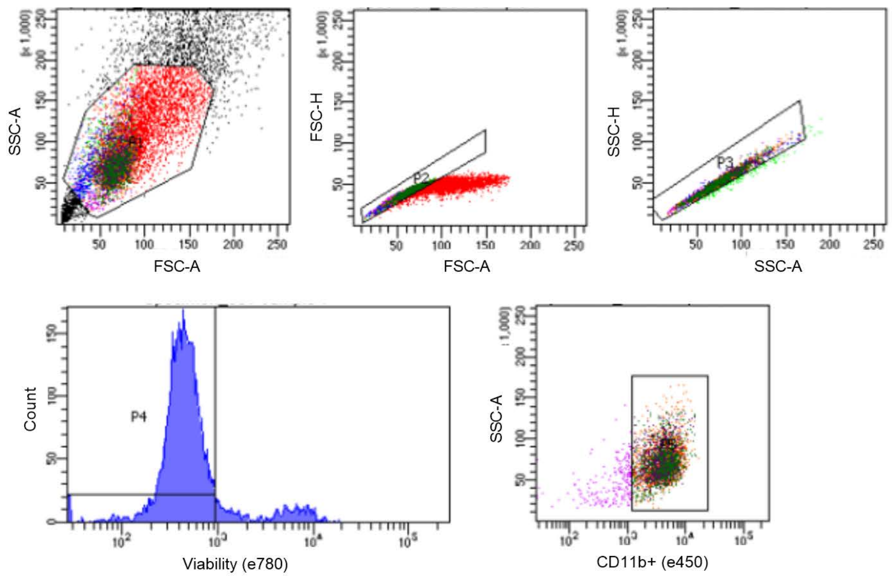


Tube: sample 1

| Population | #Events | %Parent | %Total |
|---|---|---|---|
| All Events | 13,478 | #### | 100.0 |
| P1 | 9,123 | 67.7 | 67.7 |
| P2 | 4,958 | 54.3 | 36.8 |
| P3 | 4,720 | 95.2 | 35.0 |
| P4 | 4,127 | 87.4 | 30.6 |
| P5 | 3,907 | 94.7 | 29.0 |
| PE pos | 1,218 | 31.2 | 9.0 |
| PE neg | 707 | 18.1 | 5.2 |

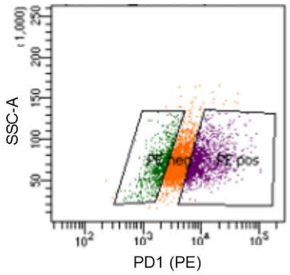


B

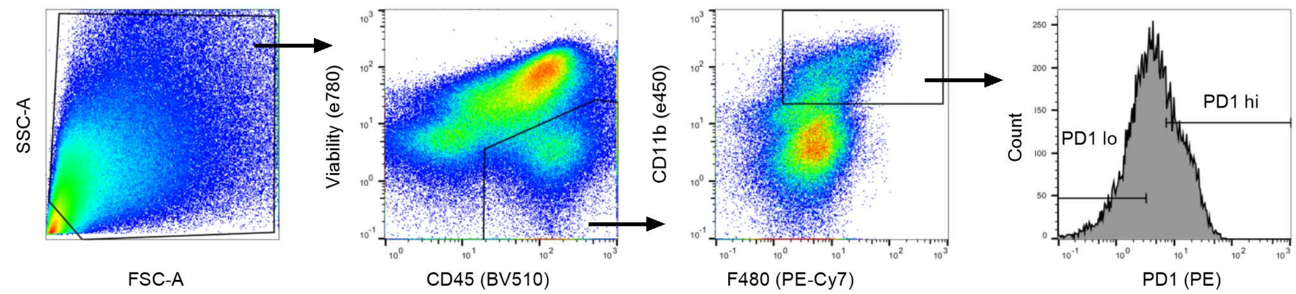

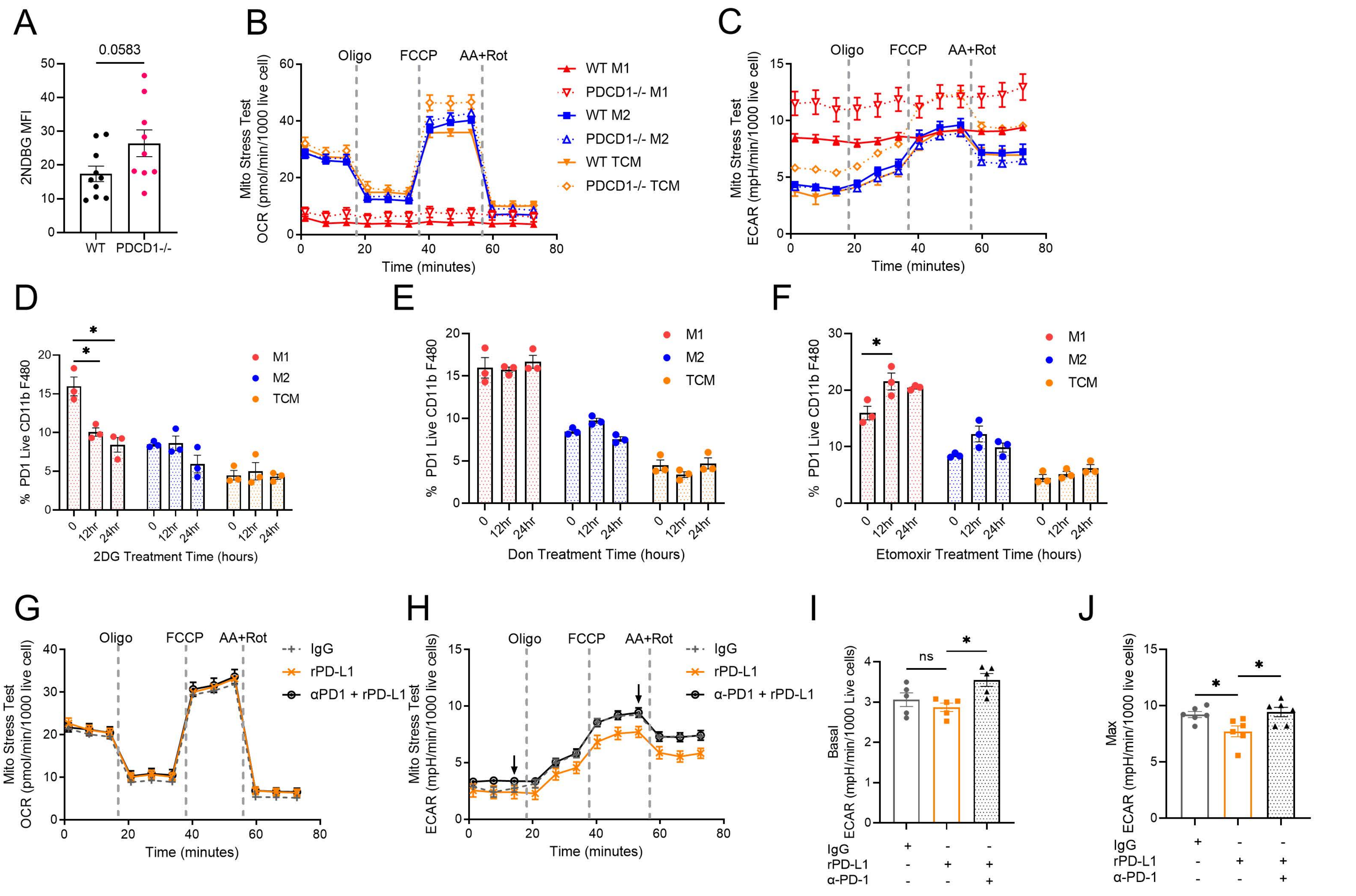

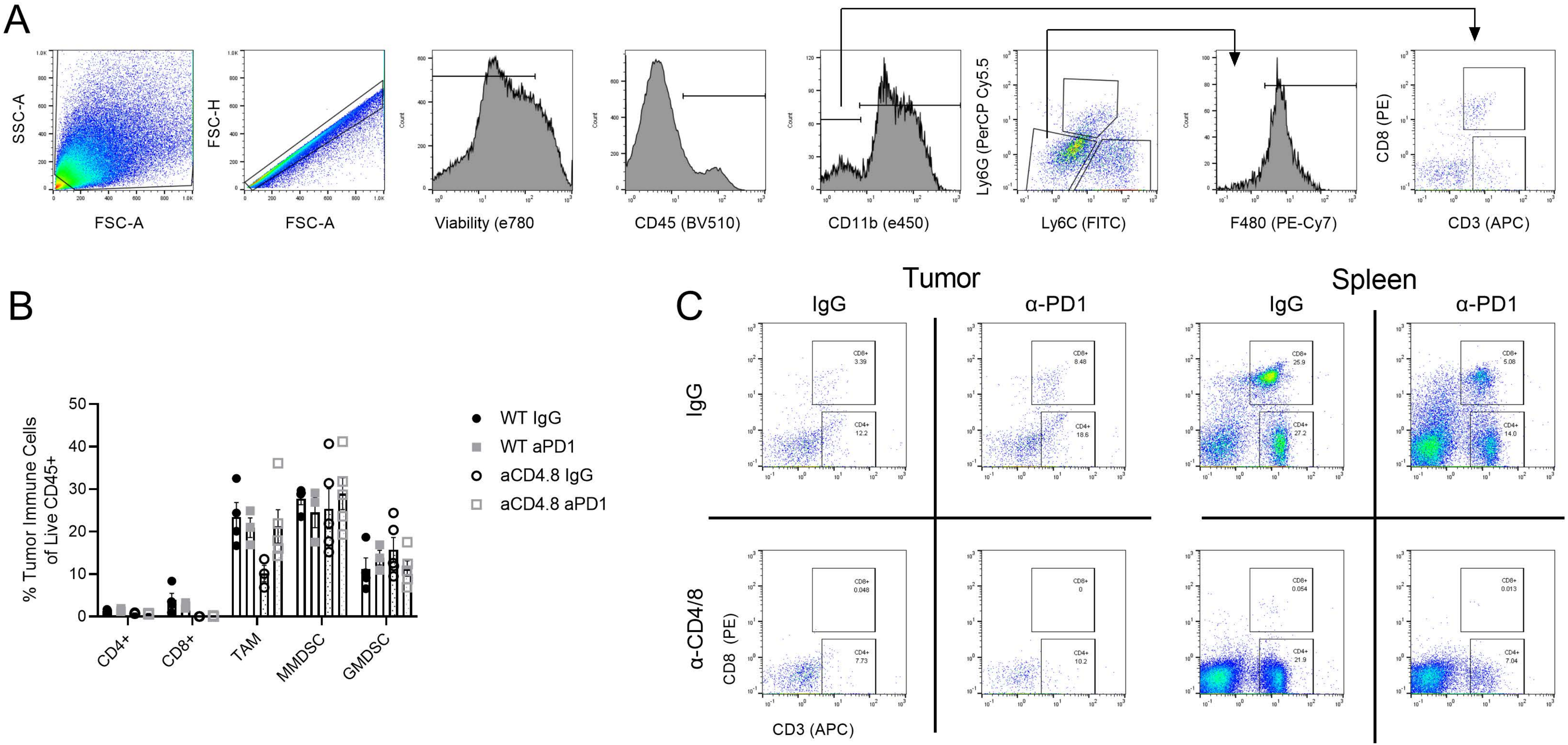

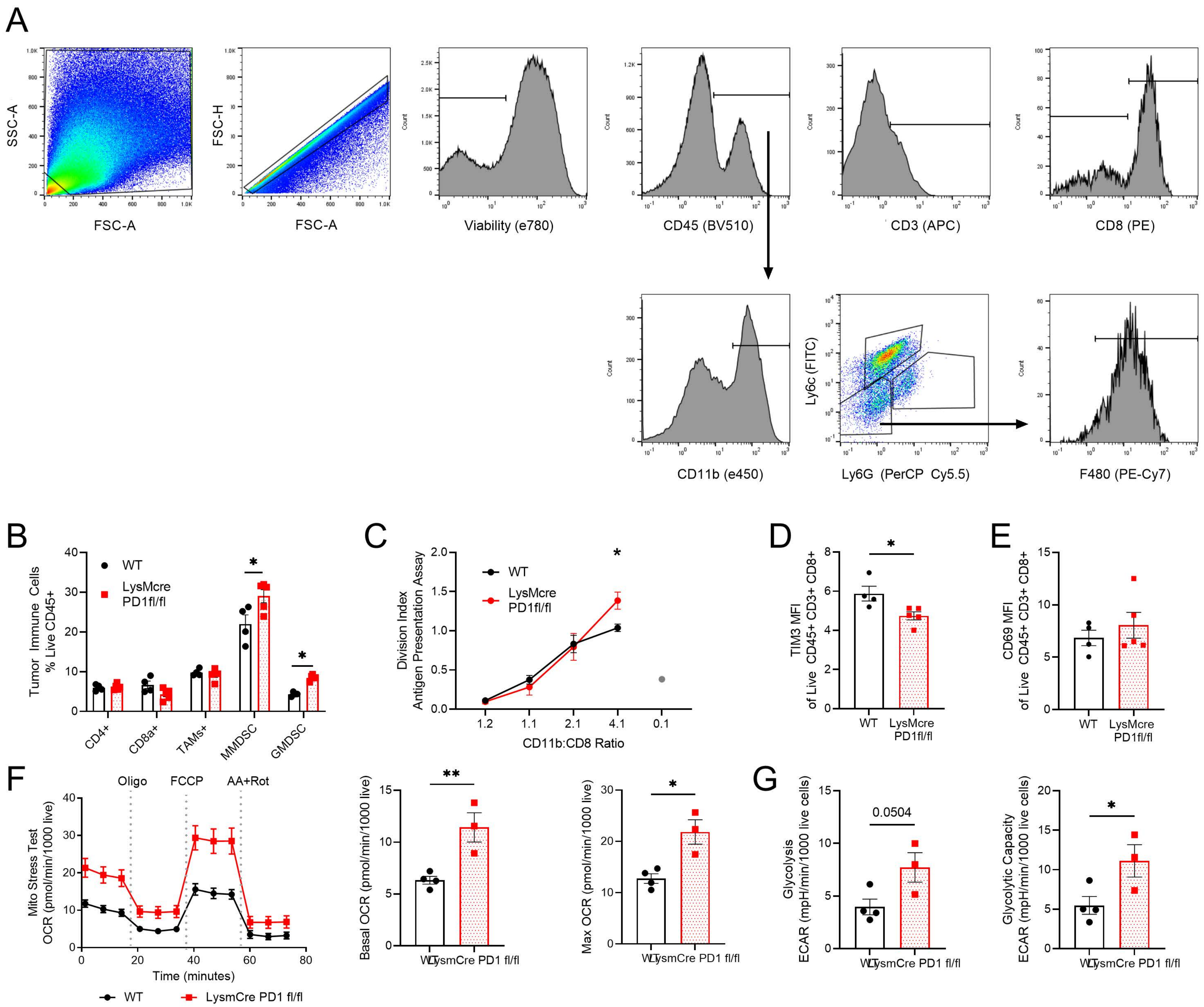
